# Foundation cell segmentation models performance on live microscopy and spatial-omics data

**DOI:** 10.64898/2026.04.18.719315

**Authors:** Yang Miao, Nick Surguladze, Josh Lerner, Koravit Poysungnoen, Ky Ariano, Yuexi Li, Yining Zhu, Kyra Van Batavia, Jodie Jepson, Josie Van De Klashorst, Bobby Y.X. Ni, Alexander Armstrong, Reeha Rahman, Roarke Horstmeyer, John W. Hickey

## Abstract

Accurate cell segmentation is an essential step for quantitative analysis of biological imaging data. Recent advances in deep learning have led to the development of generalist segmentation models that perform robustly across multiple imaging modalities, including label-free phase contrast, fluorescence cell culture, and multiplexed fluorescence tissue imaging. However, systematic comparisons of these models at the level of downstream biological analysis remain limited. To address this gap, we evaluated several recent segmentation models, including Cellpose cyto3, Cellpose-SAM, µSAM, and CellSAM, on phase contrast and fluorescence cell culture images. In addition, Mesmer and InstanSeg were included for benchmarking on multiplexed fluorescence tissue images generated using CO-Detection by IndEXing (CODEX). We found that Cellpose-SAM achieved strong performance on phase contrast images, while SAM-based models consistently performed well on fluorescence cell culture data. In contrast, no single model consistently outperformed others on CODEX datasets. Instead, each model exhibited distinct strengths and limitations, which led to differences in downstream analyses, including clustering and cell type identification. Together, our study emphasizes the importance of selecting segmentation models based on dataset characteristics and analytical goals, rather than relying on a single universal approach.

## Introduction

Imaging-based single-cell analysis has transformed biology by enabling the systematic study of cellular morphology, molecular state, and spatial organization. In cultured cells, high-throughput live-cell microscopy approaches such as cell painting^1^ and phase contrast imaging^2^ enable large-scale characterization of dynamic cellular phenotypes. In tissues, spatially resolved technologies, including spatial proteomics and spatial transcriptomics, allow the mapping of cellular composition and interactions within intact biological contexts^3–5^. These imaging modalities provide opportunities to quantify cellular behaviors and tissue organization at single-cell resolution.

A fundamental prerequisite for extracting quantitative information from these imaging datasets is accurate cell segmentation. Cell segmentation directly affects nearly all downstream analyses, including morphology quantification, feature extraction, and cell type classification. However, despite decades of algorithmic development, robust cell segmentation remains challenging due to variations in cell morphology, imaging modality, signal-to-noise ratios, and complex cellular environments, particularly in dense tissues.

Recent advances in deep learning have largely improved cell segmentation performance. Generalist segmentation models such as Cellpose^6^ and its derivatives^7–9^ provide robust performance across diverse imaging modalities and serve as strong base models without requiring extensive user-specific training. In parallel, models specifically designed for fluorescence tissue images, such as Mesmer^10^, have been widely adopted in spatial proteomics applications. Other efforts, such as InstanSeg^11^, aim to improve computational efficiency for large-scale tissue imaging datasets. More recently, foundation-model-based approaches leveraging the Segment Anything Model (SAM) framework^12^ have been introduced, including methods such as Cellpose-SAM^9^, µSAM^13^, and CellSAM^14^. These models aim to generalize across a wide range of biological images by combining large-scale pretraining with specialized decoding strategies for cellular structures.

While these approaches show promise, systematic comparisons between these emerging models remain limited. Existing benchmarking studies have primarily evaluated performance on public datasets, many of which overlap with training data, potentially introducing bias. In addition, current comparisons largely rely on mask-level metrics (e.g., intersection-over-union (IoU), average precision (AP)^6,15^, F1 score, etc.), which quantify agreement with human-annotated ground truth. Although these metrics are valuable for algorithm development, they do not necessarily capture the impact of segmentation quality on downstream biological analyses. In many applications, particularly in spatial omics, the ultimate goal of segmentation is to enable accurate quantification of cellular phenotypes, protein expression, and cell type identity. Therefore, evaluating segmentation algorithms solely based on mask overlap metrics may overlook important differences in their biological utility.

To address these limitations, we systematically evaluated several recent deep learning–based generalist segmentation models across multiple imaging contexts (**Fig. 1**). Specifically, we compared Cellpose-SAM, µSAM, and CellSAM together with the widely used Cellpose cyto3 model on self-generated expert-annotated datasets from phase contrast and fluorescence imaging of cultured cells. In addition, we evaluated Mesmer and InstanSeg alongside these models on multiplexed fluorescence imaging datasets of human intestine tissue generated using spatial proteomics technologies. For multiplexed tissue imaging datasets, we extended evaluation beyond mask-level accuracy by examining the downstream impact of segmentation on biological analyses, particularly cell type classification. This dual evaluation framework enables a more comprehensive assessment of segmentation performance in realistic biological analysis pipelines.

**Figure 1.**
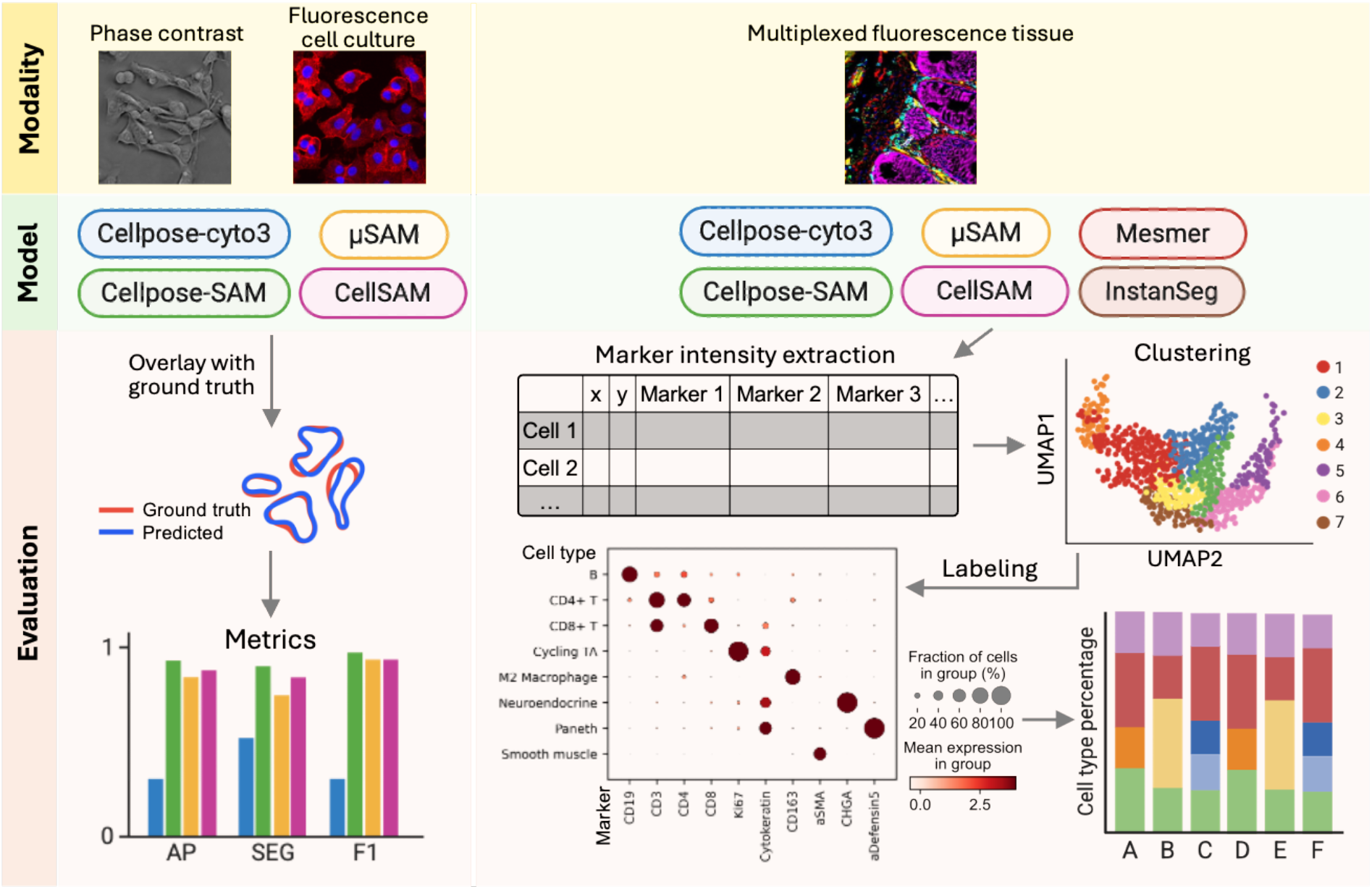
Comparison framework for segmentation models across imaging modalities. Cellpose cyto3, Cellpose-SAM, µSAM, and CellSAM were evaluated on phase contrast and fluorescence cell culture images using in-house datasets, with performance assessed by three quantitative metrics. For multiplexed fluorescence tissue images (CODEX), Mesmer and InstanSeg were additionally included. Marker intensities extracted from segmentation masks were used for downstream clustering and cell type annotation to assess the impact on biological interpretation. AP, average precision; SEG, segmentation accuracy.

## Results

### Cellpose-SAM outperforms other models on live-cell phase imaging, with additional improvements from custom training

We first assessed Cellpose cyto3, Cellpose-SAM, µSAM, and CellSAM on self-generated live-cell phase contrast images of cultured mouse melanoma B16-F10 cells. Phase contrast imaging is a label-free microscopy method that visualizes cells based on asymmetric illumination, which highlights intrinsic differences in cellular density and structure, allowing cell morphology to be observed without fluorescent staining. Because these images capture gradual variations within cells rather than sharp boundaries defined by fluorescent markers, accurately identifying individual cell borders can be challenging for segmentation algorithms. Visual inspection indicated that Cellpose-SAM successfully segmented substantially more cells than the other models, although it still missed a small number of cells (**Fig. S1A**).

To further quantify model performance, we applied the same comparison to phase contrast images of cultured human lung carcinoma A549 cells. Using expert-annotated masks as ground truth, we compared the number of predicted masks and the mean mask area produced by each model. Both Cellpose-SAM and CellSAM generated mask counts that were similar to the human annotations (**Fig. 2A**). However, CellSAM produced substantially smaller mean mask areas, suggesting that many predicted masks corresponded to partial detections (**Fig. 2B**).

**Figure 2.**
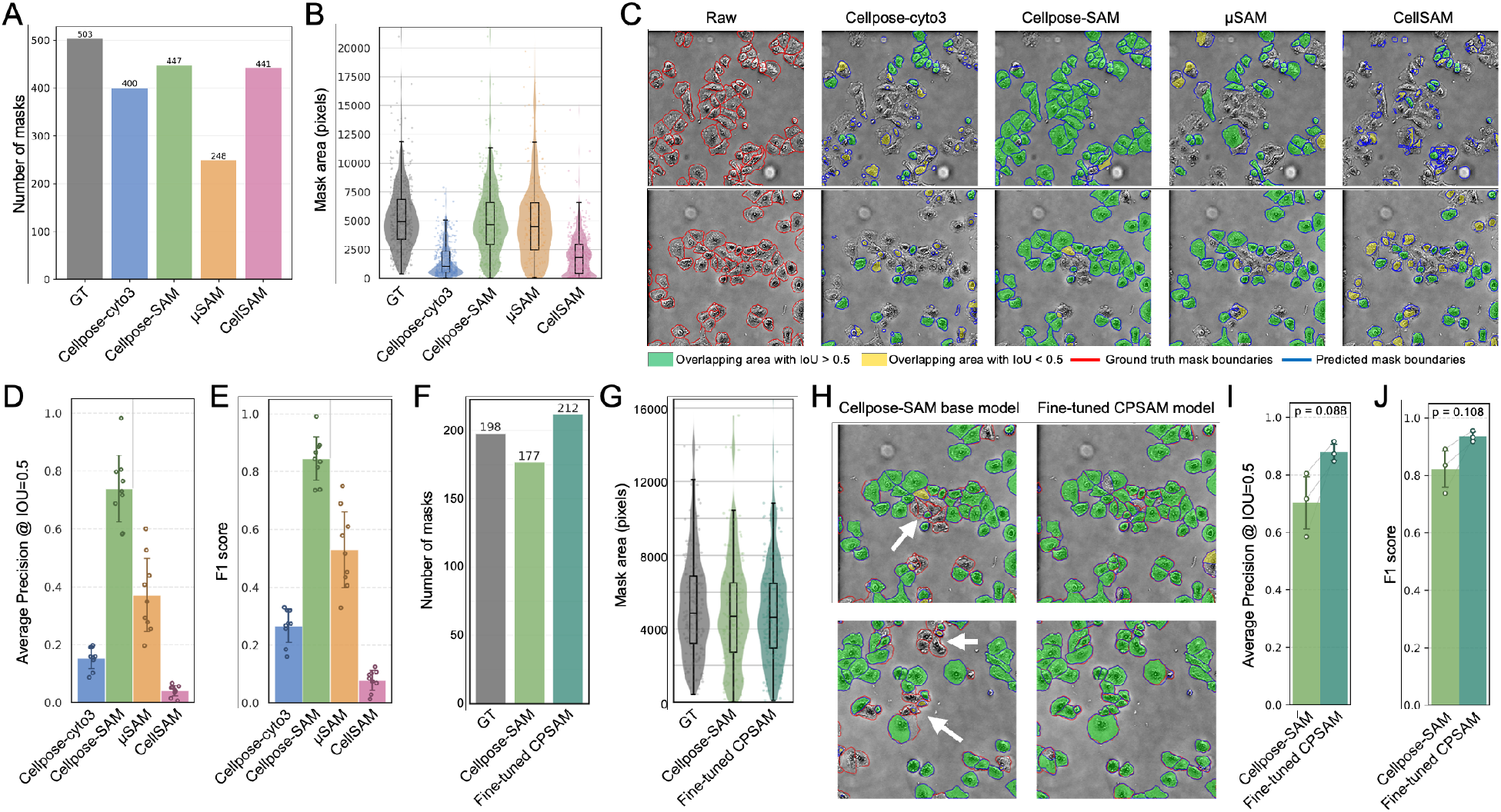
Model comparison on phase contrast images. (A) Total number of masks generated across all images. GT, ground truth. (B) Mean mask area per model. (C) Representative overlays of segmentation masks on raw images. Ground truth boundaries are shown in red and model predictions in blue. Masks matched to ground truth with IoU > 0.5 are shown in green, and true positives with IoU < 0.5 in yellow. (D) Average precision. (E) F1 score. (F) Total number of masks for Cellpose-SAM (CPSAM) and fine-tuned CPSAM. (G) Mean mask area. (H) Representative fields of view comparing base and fine-tuned CPSAM, with improved regions indicated by white arrows. (I) Average precision. (J) F1 score. P values were calculated using paired t-tests.

Visual inspection of the segmentation overlays supported these observations (**Fig. 2C**). CellSAM generated a large number of partial masks across the image. In contrast, Cellpose cyto3 and µSAM produced a higher number of accurate predictions, but they failed to segment many cells present in the image. Cellpose-SAM provided the best balance between sensitivity and specificity, successfully segmenting the majority of cells while maintaining relatively low artifactual segmentation masks, although several cells were still missed.

To quantitatively compare performance, we calculated AP, F1 score, and Segmentation Accuracy (SEG)^16^ for all models (**Methods, Fig. 2D, E, Fig. S1B**). Consistent with the visual assessment, Cellpose-SAM achieved the highest scores across all metrics, µSAM and Cellpose cyto3 showed intermediate performance, and CellSAM achieved lower scores on this dataset. Differences could come from differences in model architecture where suboptimal localization of cells for prompts could result in incomplete masks generated by SAM. Moreover, the phase contrast datasets used for model training may differ from our datasets, which could further limit model generalization. As a result, models may fail to accurately detect cells in imaging data with distinct modality-specific characteristics and contrast features.

Although the base Cellpose-SAM model achieved the best overall performance, it still failed to detect several cells. To test whether model adaptation could further improve performance, we performed custom training of Cellpose-SAM using a subset of expert-annotated masks and human-in-the-loop retraining^7^, while reserving the remaining annotations for validation. The fine-tuned model predicted a slightly larger number of cells than expected (**Fig. 2F**) with similar size (**Fig. 2G**), but inspection of the overlays showed that it successfully recovered several cells missed by the base model (**Fig. 2H**). Quantitative evaluation indicated consistently higher segmentation metrics compared to the pretrained model, although the improvement did not reach statistical significance (**Fig. 2I, J, Fig. S1C**). Together, these results demonstrate that Cellpose-SAM provides the most accurate segmentation among the tested generalist models for phase imaging of cultured cells, and that improvements can be achieved through dataset-specific fine-tuning (**Table 1**).

**Table 1.** Summary of segmentation performance across models on phase contrast and fluorescence cell culture images.

| | | Cellpose cyto3 | Cellpose-SAM | μSAM | CellSAM |
| --- | --- | --- | --- | --- | --- |
| Phase contrast | General performance | * | *** | ** | * |
| Fluorescence cell culture | General performance | ** | *** | *** | *** |
|  | Low S/N ratio robustness | * | *** | *** | *** |

\*\*\* Comparable to human expert performance
\*\* Moderate performance
\* Lower performance

### SAM-based models achieve high segmentation accuracy on fluorescence images of cultured cells and show strong robustness to signal-to-noise variation

We next evaluated the three SAM-based models, Cellpose-SAM, µSAM, and CellSAM together with the Cellpose cyto3 model on fluorescence images of fixed A549 cells. Fluorescence imaging is widely used in biological research because it enables specific labeling of cellular structures, allowing precise visualization and quantitative analysis of cell morphology and organization. In these images, cells were labeled with a cytoplasmic protein marker (red) and a nuclear marker (blue) captured in separate fluorescence channels, and the two channels were provided separately to each model according to their recommended channel configuration.

All three SAM-based models produced a number of predicted masks that closely matched the human-annotated ground truth (**Fig. 3A**), while Cellpose cyto3 produced masks with substantially smaller average areas (**Fig. 3B**). Visual inspection of segmentation overlays showed that the SAM-based models accurately captured full cell morphology, including cytoplasmic boundaries. In contrast, Cellpose cyto3 frequently failed to recover complete cell shapes and detected mainly the nuclei, likely due to the strong and clear nuclear signal present in the images. As a result, while Cellpose cyto3 could identify the presence of cells, it often underestimated the full cellular boundaries (**Fig. 3C**). Consistent with visual inspection, all three SAM-based models achieved accurate AP, F1 score and SEG scores that were comparable to expert human annotations with Cellpose cyto3 having lower performance and greater variation (**Fig. 3D, E, Fig. S2A**).

**Figure 3.**
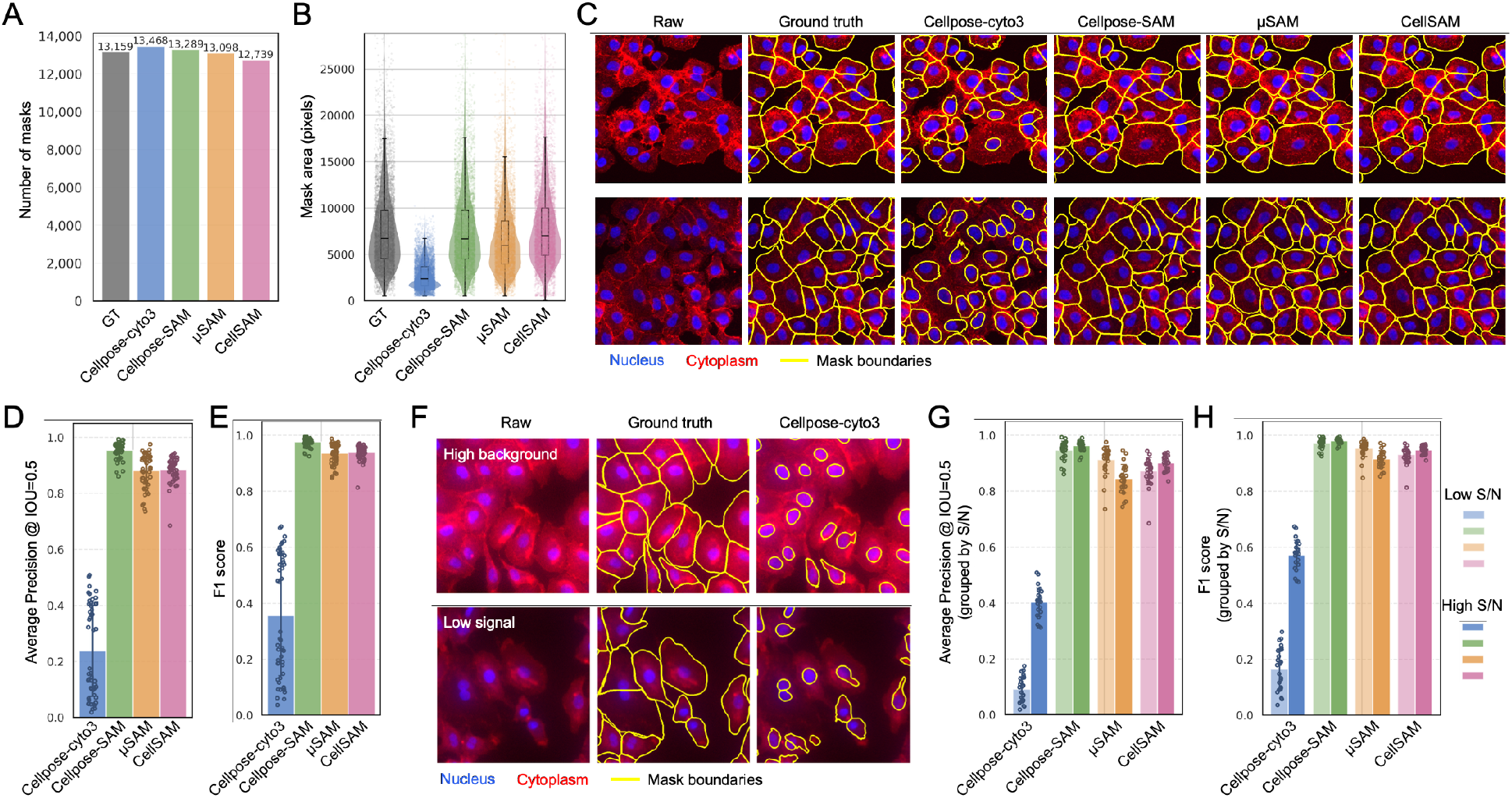
Model comparison on fluorescence cell culture images. (A) Total number of masks generated across all images. GT, ground truth. (B) Mean mask area per model. (C) Representative overlays of segmentation masks on raw images. Nucleus marker is shown in blue and membrane marker in red; mask boundaries are shown in yellow. (D) Average precision. (E) F1 score. (F) Representative low signal-to-noise (S/N) examples, including high background and low signal conditions, showing Cellpose cyto3 segmentation. (G) Average precision grouped by high and low S/N conditions. (H) F1 score grouped by high and low S/N conditions.

We hypothesized that this variability in Cellpose cyto3 may be driven by differences in fluorescence image signal-to-noise (S/N) ratio. To test this, we divided the dataset into two groups: high S/N images and low S/N images. Examples of low S/N images included cases with high background fluorescence and cases with weak staining resulting in low signal intensity. Both scenarios commonly encountered in biological experiments. In these challenging conditions, Cellpose cyto3 typically detected nuclei but failed to reconstruct full cell boundaries (**Fig. 3F**). In contrast, all three SAM-based models maintained accurate segmentation of cell morphology (**Fig. S2B**).

Quantitative analysis supported these observations. Cellpose cyto3 performance improved substantially when moving from low S/N images to high S/N images (**Fig. 3G, H, Fig. S2C**). This result indicates a strong dependence on input image quality for Cellpose cyto3. In comparison, the three SAM-based models maintained consistently high performance across both conditions, demonstrating greater robustness to variations in fluorescence signal quality (**Table 1**).

### Segmentation models exhibit distinct strengths on multiplexed fluorescence tissue images, with SAM-based models producing the most similar predictions

We next evaluated segmentation performance on multiplexed fluorescence spatial proteomics images. Multiplexed fluorescence imaging enables simultaneous measurement of dozens of protein markers within the same tissue section, allowing high-dimensional characterization of cellular phenotypes and spatial organization in complex tissues. In such datasets, protein marker intensities are typically quantified from segmented cell masks to generate feature vectors that are subsequently used for clustering and cell type annotation.

We selected eight human large and small intestine tissue images from the Human Biomolecular Atlas Program (HuBMAP) generated using the Co-Detection by Indexing (CODEX) spatial proteomics assay^5,17^. These datasets contain more than 40 protein marker fluorescence channels. For segmentation input, we used the Hoechst channel as the nuclear signal and a composite of Cytokeratin (epithelial marker) and CD45 (immune marker) as the whole-cell or membrane channel. As a reference for comparison, we used the consortium-provided segmentation masks generated using the Cytokit pipeline^18^. In addition to Cellpose cyto3, Cellpose-SAM, µSAM, and CellSAM, we also included Mesmer and InstanSeg, two models that have been specifically adapted for fluorescence tissue imaging.

Visual inspection across representative tissue regions, including epithelial compartments, stromal regions, and immune follicle areas, revealed distinct strengths across models (**Fig. 4A**). In intestinal epithelial regions, where cells exhibit densely packed columnar morphology, Cellpose-SAM produced particularly accurate segmentation boundaries. Mesmer and Cellpose cyto3 also achieved relatively high accuracy in segmenting these epithelial cells. However, epithelial staining quality varied across regions, and we observed that the advantage of Cellpose-SAM in resolving epithelial cells was most evident when staining signals were strong and well-defined. Under suboptimal staining conditions, where signals were weak or less distinct, this advantage was reduced, and Cellpose-SAM no longer consistently outperformed the other models (**Fig. S3A**).

**Figure 4.**
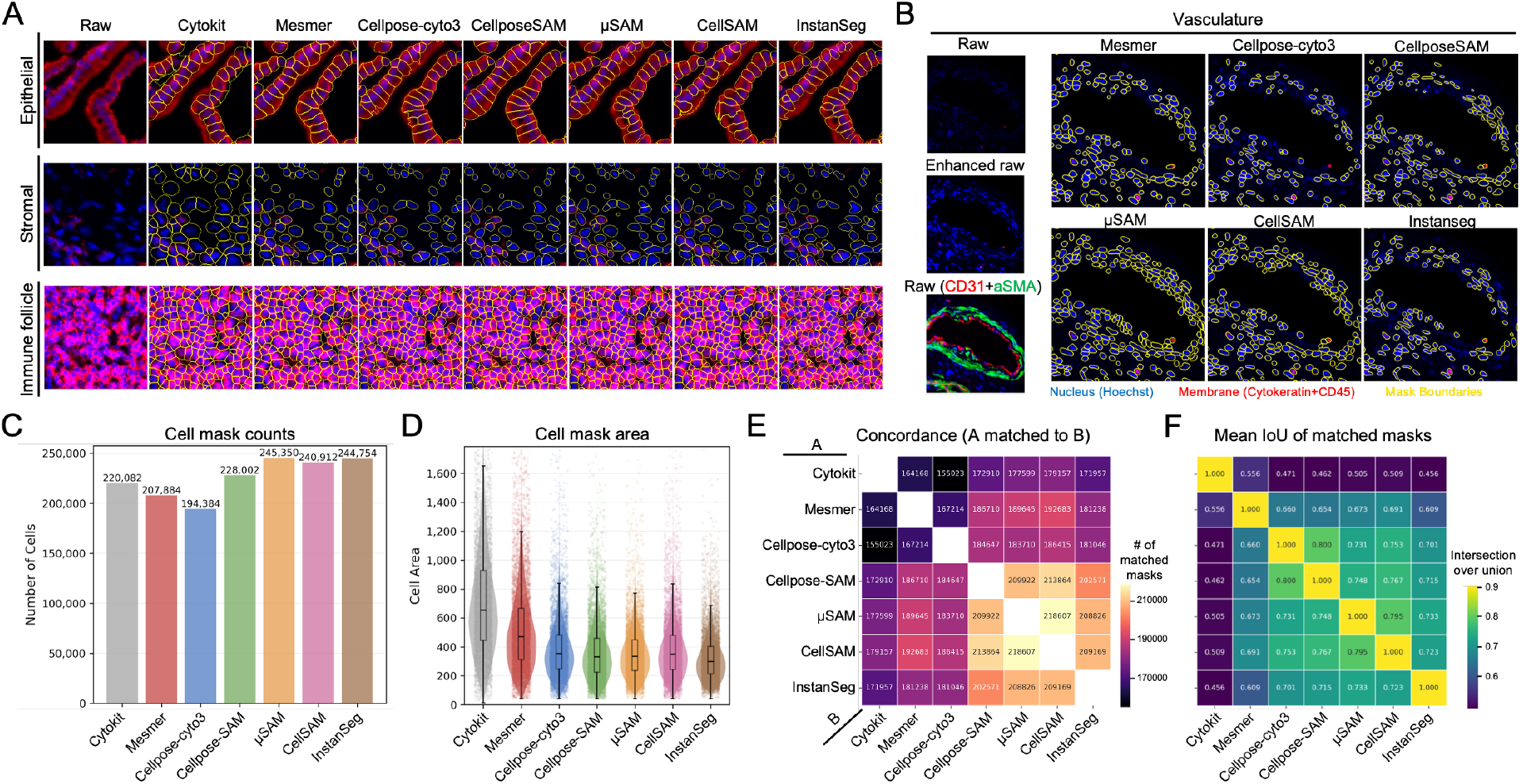
Model comparison on multiplexed fluorescence CODEX tissue images. (A) Representative overlays of segmentation masks on epithelial, stromal, and immune follicle regions. Nucleus is shown in blue, membrane in red, and mask boundaries in yellow. (B) Representative vasculature region with segmentation overlays. Raw images without contrast enhancement show weak nuclear signal, whereas enhanced images improve nuclear visualization. CD31 (red) and αSMA (green) highlight vascular structure. (C) Total number of masks generated across all images. (D) Mean mask area per model. (E) Pairwise mask matching between models; colors indicate the number of matched masks. (F) Mean IoU of matched masks in (E); colors indicate average IoU values.

In stromal and immune follicle regions, most models produced similar predictions, likely because cells in these regions exhibit clearer nuclear and membrane signals that simplify segmentation.

In contrast, we observed more visual differences in segmentation of vascular-associated regions (**Fig. 4B**). Cells surrounding vasculature often exhibit heterogeneous morphology and variable nuclear shapes, and their nuclei can be substantially dimmer than cells in other areas. These characteristics make segmentation difficult for many algorithms. In these regions, µSAM and CellSAM demonstrated higher sensitivity to weak nuclear signals and successfully detected nearly all cells present in the area. Mesmer showed moderate performance, while Cellpose cyto3, Cellpose-SAM, and InstanSeg missed a larger number of cells with dim nuclei.

We also observed differences in segmentation robustness in regions with blurred fluorescence signals, which are common in multiplexed spatial proteomics datasets due to optical limitations or imaging artifacts. Mesmer and µSAM appeared particularly robust in these regions and were able to successfully segment cells even when signal boundaries were blurred. CellSAM and InstanSeg also showed relatively good performance, while Cellpose-SAM and Cellpose cyto3 were more sensitive to signal blur and often failed to recover complete cell masks in these regions (**Fig. S3B**).

To quantify these observations, we compared the number and area of predicted cell masks across models. µSAM, CellSAM, and InstanSeg produced the largest number of cell masks, while Cellpose cyto3 predicted the fewest cells (**Fig. 4C**). In particular, Cellpose cyto3 frequently failed to segment cells in specific tissue regions, especially in smooth muscle and stromal areas that have lower signal to noise ratio nuclei (**Fig. S3C**). We also observed that µSAM occasionally over-segmented regions by detecting weak nuclear signals that likely correspond to noise, though these segmentation artifacts can potentially be filtered during downstream analysis by applying thresholds on nuclear signal intensity. In terms of mask size, Cytokit produced the largest average cell mask areas, consistent with visual inspection of its segmentation results, followed by Mesmer (**Fig. 4D**).

We further examined the similarity of segmentation predictions between models by implementing a cell-mask matching strategy. In this approach, two masks from a model pair were considered matched if each mask was the nearest neighbor of the other (mutual nearest neighbors) based on the distance between their centroids, and if this distance was below a predefined spatial threshold (**Methods**). Interestingly, the three SAM-based models, Cellpose-SAM, µSAM, and CellSAM, showed the highest degree of agreement with each other. This similarity likely arises from their shared foundation in the SAM architecture, which preserves common feature representations across models (**Fig. 4E**).

We also quantified the mean Intersection-over-Union (IoU) of matched cell masks between each pair of models. Among matched cells, Cellpose cyto3 and Cellpose-SAM produced the most similar mask shapes (**Fig. 4F**). This result is consistent with the design of Cellpose-SAM, which integrates SAM-derived image features into the Cellpose framework while retaining the same flow-based mask reconstruction strategy. As a result, Cellpose-SAM preserves the shape priors of the original Cellpose model, leading to similar segmentation geometries. The next highest similarity was observed between µSAM and CellSAM, which both adopt a prompt-based segmentation strategy in which intermediate model components generate prompts that are subsequently passed to the SAM prompt encoder and mask decoder to produce final cell masks. These observations suggest that differences in model architecture influence cell mask generation. Such differences become particularly obvious in challenging tissue regions, including densely packed areas and regions with weak or ambiguous fluorescence signals.

Another important consideration when applying segmentation models to multiplexed fluorescence imaging datasets is computational efficiency. These datasets are typically very large and therefore require substantial computational resources and processing time. To benchmark runtime performance, we processed eight CODEX tissue regions (∼10,000 × 10,000 pixels per image, corresponding to ∼3.77 mm × 3.77 mm tissue area) using an NVIDIA RTX 4080 GPU.

Among all tested models, CellSAM required the longest processing time, taking approximately 8 hours in total (∼60 minutes per region). This longer runtime is partly due to our use of a relatively small tile size (250 pixels) to improve segmentation accuracy after multiple rounds of parameter tuning. In addition, we enabled the *gauge_cell_size* option, which automatically estimates cell size and further increases computational cost. The second most computationally intensive model was µSAM, which required 33 minutes total (∼4 minutes per region) when using the AutomaticPromptGenerator framework with the heavier ViT-L backbone model (vit_l_lm), a tile size of 512 pixels, and parameters optimized for segmentation performance. Mesmer required 25 minutes total (∼3 minutes per region), while Cellpose-SAM processed the dataset in 16 minutes (∼2 minutes per region). InstanSeg and Cellpose cyto3 were the most computationally efficient methods, requiring approximately 6.5 minutes (∼47 seconds per region) and 6 minutes (∼44 seconds per region), respectively (**Table 2**). These results highlight substantial differences in computational cost among segmentation models when applied to large spatial proteomics datasets. While some models achieve improved segmentation sensitivity or robustness, they may require longer processing times, which becomes an important practical consideration for large-scale spatial omics studies (**Table 2**).

**Table 2.** Summary of mask-level segmentation performance across models on multiplexed fluorescence (CODEX) images.

| | Mesmer | Cellpose-cyto3 | Cellpose-SAM | μSAM | CellSAM | InstanSeg |
| --- | --- | --- | --- | --- | --- | --- |
| **Mask level** |  |  |  |  |  |  |
| Mask counts | 207884 | 194384 | 228002 | 245350 | 240912 | 244754 |
| Mask area (pixels, mean ± SD) | 525 ± 301 | 396 ± 226 | 371 ± 218 | 365 ± 193 | 390 ± 221 | 331 ± 169 |
| Epithelial compartment performance | * | * | ** | * | * | * |
| Stromal compartment performance | *** | *** | *** | *** | *** | *** |
| Follicle compartment performance | *** | *** | *** | *** | *** | *** |
| Dim nucleus sensitivity | ** | * | ** | *** | *** | * |
| Signal blur robustness | *** | * | * | *** | *** | ** |
| Background false positives | No | No | No | Yes | No | No |
| Computation time (per sample) | 3 min | 44 sec | 2 min | 4 min | 60 min | 47 sec |

\*\*\* Comparable to human expert performance
\*\* Moderate performance
\* Lower performance

### Segmentation differences propagate into downstream biological analyses and affect cell type resolution

Given the substantial differences in segmentation masks observed across models on multiplexed fluorescence images, we next asked whether these differences would impact downstream biological analyses. However, most existing studies focus primarily on mask-level performance metrics or qualitative visual inspection, and do not systematically evaluate how segmentation variability affects downstream readouts such as cell type composition. As a result, the extent to which segmentation choice influences biological conclusions remains unclear. Although some prior work has examined the ability of individual methods, such as Mesmer, to identify specific cell types, comprehensive comparisons across multiple state-of-the-art segmentation models in downstream analysis tasks are still lacking.

To evaluate this, we applied our standard CODEX data processing pipeline to each segmentation output^17,19^. Briefly, we extracted the mean fluorescence intensity of each marker channel within every cell mask to generate a single-cell expression vector. Marker intensities were then z-normalized across cells for each channel and removed problematic masks based on size and signal saturation (**Methods**). After filtering, we performed unsupervised Leiden clustering using the same resolution parameters and a predefined marker panel to identify epithelial, stromal, and immune cell populations (**Fig. 5A**). The resulting clusters were manually inspected and annotated based on their marker expression patterns. This entire procedure was repeated for each segmentation model. As a reference, we used human expert annotations from the HuBMAP consortium generated from Cytokit-based segmentation of the same datasets, which followed a similar annotation workflow.

**Figure 5.**
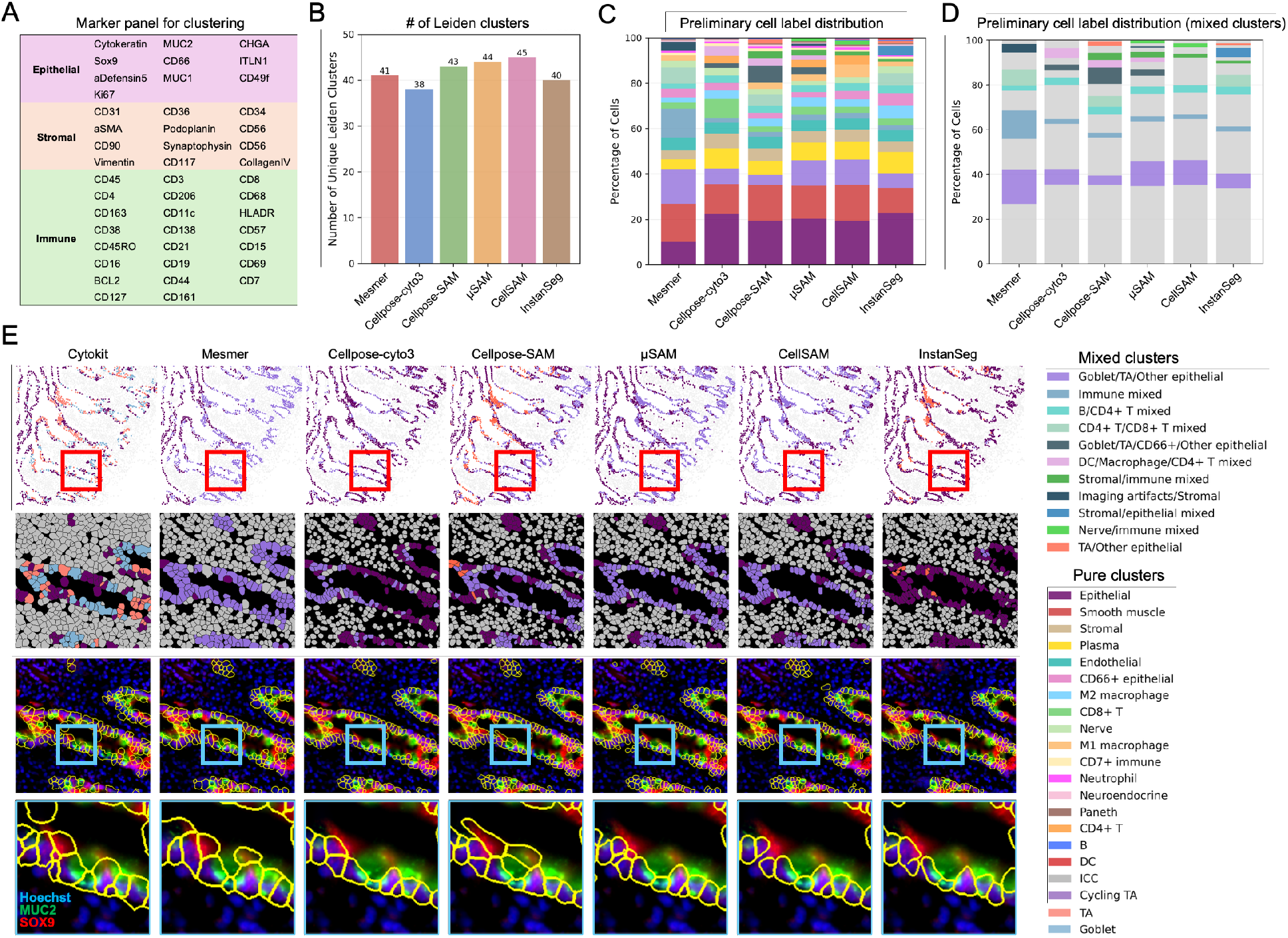
Downstream cell type classification across segmentation models on CODEX images. (A) Marker panel used for clustering, highlighting epithelial, stromal, and immune cell types. (B) Number of unsupervised Leiden clusters per model. (C) Cell type proportions after manual annotation of Leiden clusters and merging of clusters with identical labels. (D) Proportion of mixed clusters only. (E) Representative tissue region illustrating differences in epithelial segmentation. Top row, overview region; red box indicates the area shown below. Second row, segmentation masks with epithelial-containing clusters colored. Third row, mask boundaries overlaid on raw images. Bottom row, zoomed-in region (blue box) highlighting local differences in marker capture across models. Nucleus is shown in blue, goblet marker MUC2 in green, and TA marker SOX9 in red. Cell type color legend is shown on the right.

Across all segmentation models, Leiden clustering produced the highest number of clusters in CellSAM and the fewest in Cellpose cyto3, suggesting differences in how well cell populations were separated based on their marker expression profiles (**Fig. 5B**). To assess cluster quality, we manually annotated each Leiden cluster for each model by examining marker expression patterns across cells within each cluster and visually confirming these patterns against the raw staining in the original images. We observed that clusters frequently contained a mixture of multiple cell types, suggesting that cells with distinct identities were often grouped together during clustering.

Based on these observations, clusters were categorized using the following criteria. Clusters in which more than 90% of cells corresponded to a single cell type were defined as pure clusters and labeled accordingly (e.g., Paneth, smooth muscle). Clusters containing multiple cell types were defined as mixed clusters and labeled by their predominant cell types based on visual inspection (e.g., CD4+ T/CD8+ T mixed, stromal/immune mixed). In total, 251 Leiden clusters were manually annotated. Clusters with identical labels were subsequently merged within each model to generate the final set of cell type assignments (**Supplemental Files 1–7**).

In total, 20 pure cell types were identified, of which 14 were consistently observed across all models, while the remaining cell types appeared as pure clusters only in a subset of models (**Fig. 5C**). In addition, 11 mixed cluster types were identified, among which only three (B/CD4+ T mixed, goblet/TA/other epithelial, and immune mixed) were consistently observed across all models, whereas the remaining eight were specific to one or multiple models (**Fig. 5C, D**).

Within the pure clusters, segmentation differences influenced the ability to resolve certain biologically important cell populations. Notably, only Mesmer identified a distinct cluster corresponding to interstitial cells of Cajal (ICC), characterized by co-expression of CD44 and CD117 (**Table S1, Supplemental File 1**). ICCs are relatively rare cells located in the muscularis propria layer of the intestine and play an important role in regulating intestinal motility. Mesmer’s ability to capture this population may be related to its larger segmentation masks, which capture more surrounding fluorescence signal and may enhance detection of weak marker combinations. Conversely, only Cellpose-SAM and CellSAM identified a distinct cluster of cycling transit-amplifying (Cycling TA) epithelial cells (**Table S1, Supplemental File 3, 5**), which is a cell type that typically resides near the base of intestinal crypts and play a key role in epithelial renewal, marked by strong Ki67 expression together with epithelial markers such as Cytokeratin. In addition, only CellSAM and InstanSeg resolved a distinct cluster of dendritic cells (**Table S1, Supplemental File 5, 6**), marked by CD11c and HLA-DR expression. Dendritic cells are key antigen-presenting cells in the adaptive immune response but often exhibit heterogeneous morphology, making them difficult to segment accurately and prone to being missed or misclassified. By contrast, none of the segmentation models identified a distinct cluster corresponding to natural killer (NK) cells, despite their presence in HuBMAP expert annotations (**Table S1**). This may be due to the low abundance of NK cells in the tissue and the limited specificity of commonly used markers, such as CD56 and CD57, which can overlap with other cell populations and hinder reliable cluster separation.

Beyond pure clusters, we observed 11 clusters containing mixed cell populations that varied in proportion across segmentation models (**Fig. 5D, Fig. S4A**). Cellpose cyto3 and CellSAM produced the lowest overall proportion of mixed-cell clusters, although the specific cell types involved differed between the models. Mesmer showed the highest overall proportion of mixed clusters. This may be explained by its tendency to generate larger segmentation masks, which can capture fluorescence signals from adjacent cells and lead to signal contamination. Interestingly, Mesmer did not produce a larger number of mixed clusters overall, suggesting that certain cell populations may be more prone to mixing rather than increasing the diversity of mixed populations. Cellpose-SAM and µSAM produced the highest number of mixed clusters and also showed relatively high overall proportions of mixed-cell populations, second only to Mesmer.

To further investigate these mixed populations, we examined which cell types were most frequently grouped together. Across all models, epithelial subtypes were particularly difficult to resolve. These included goblet cells (MUC2+), transit-amplifying cells (SOX9+), and other epithelial subtypes (ITLN+, MUC1+). We selected a representative region from the small intestine to illustrate these challenges (**Fig. 5E**). In this region, Cytokeratin staining was less clear than in other epithelial areas, making cell boundaries less distinct (**Fig. S4B**).

Although none of the segmentation models perfectly resolved individual cells, including Cellpose-SAM, the masks generated by different models varied substantially. Even small segmentation differences altered the measured marker intensities for MUC2 and SOX9, leading to different clustering outcomes. Biologically, goblet cells and TA cells often reside in close proximity within intestinal crypts. Goblet cells secrete mucus, while TA cells migrate upward during epithelial renewal. Their close spatial arrangement makes signal spillover between adjacent cells common, further complicating segmentation-based quantification.

To quantify the impact of these segmentation differences, we compared the proportions of epithelial subtypes identified by each model in this region (**Fig. S4C**). Substantial variation was observed across models. In practical workflows, clusters initially labeled as “pure epithelial” without strong expression of MUC2 or SOX9 are often not further refined during manual annotation. As a result, segmentation differences at this stage can directly affect the estimated proportions of goblet cells, TA cells, and non-specialized mature epithelial cells. For example, InstanSeg assigned a high proportion of epithelial cells in this region to non-specialized mature epithelial cells (**Fig. S4C**), indicating a failure to resolve goblet and TA populations. In contrast, Mesmer classified a low proportion of epithelial cells as non-specialized mature cells, suggesting a higher likelihood of identifying goblet and TA populations during subsequent refinement. To further assess this effect, we performed k-means subclustering (k = 3) on the mixed epithelial clusters within this region to mimic a typical annotation refinement step (**Fig. S4D**). Clusters were defined based on marker expression as TA (SOX9 high), goblet (MUC2 high), and a remaining group corresponding to other epithelial cells. As expected, subclustering enabled the separation of goblet and TA populations from the mixed clusters. Among all models, Mesmer produced cell-type proportions that were most consistent with human expert annotations following this refinement step (**Fig. S4E**).

Another example of epithelial subtype mixing involved CD66+ epithelial cells, which are primarily located near the luminal surface of the large intestine. Although all segmentation models produced a cluster enriched for CD66+ epithelial cells, several models, including Cellpose cyto3, Cellpose-SAM, and µSAM, failed to capture all CD66+ epithelial cells within this cluster, indicating incomplete separation of this population (**Fig. S5A, B, D**).

Immune cell subtypes also exhibited frequent mixing during clustering. Examples included mixing between CD4+ T cells and CD8+ T cells, B cells and CD4+ T cells, and dendritic cells with macrophages or CD4+ T cells (**Fig. 5D**). Such mixing reflects both biological and technical challenges. Many immune cell types share common markers, including CD45 for all immune cells, CD3 for T cells, and HLA-DR for antigen-presenting cells. Additionally, macrophage subtypes such as M1 and M2 macrophages often differ primarily in expression levels of markers like CD68 and HLA-DR, making them difficult to distinguish computationally. Despite these challenges, CellSAM and InstanSeg showed the strongest ability to separate dendritic cells from M1/M2 macrophages (**Table S1, Supplemental File 5, 6**), whereas Mesmer failed to distinguish dendritic cells and instead grouped them within a mixed immune cluster (**Table S1, Supplemental File 1**). All models struggled to fully separate B cells from CD4+ T cells, likely due to spatial proximity in follicle areas (**Table S1**). However, CellSAM and µSAM produced relatively pure CD8+ T cell clusters compared to other models (**Table S1, Supplemental File 4, 5**).

We also observed that although all segmentation models produced a cluster labeled as immune mixed, Mesmer contained a noticeably higher fraction of plasma cells within this group (**Fig. S5C, E**). Plasma cells are abundant beneath epithelial layers where mucosal immune responses occur. Because Mesmer tends to generate larger masks, signals from plasma cell markers such as CD38 and CD138 may be diluted by signals from neighboring cells, reducing clustering specificity.

Taken together, our analyses show that all segmentation models led to different degrees of cell-type mixing in downstream clustering, both in the overall proportion of mixed clusters and in the specific cell populations affected. No single model was uniformly best across all criteria, and each model showed a different balance between clustering purity, biological resolution, and computational efficiency.

CellSAM produced the lowest overall proportion of mixed clusters, suggesting relatively strong performance in preserving cluster purity. However, this advantage came with substantially longer computational time. Mesmer showed the highest proportion of mixed cells, but the total number of mixed cluster types was similar to that of CellSAM, indicating that the practical amount of downstream refinement required by users may still be comparable. Cellpose cyto3 and InstanSeg were the most computationally efficient models, making them attractive for large-scale analysis, although Cellpose cyto3 frequently missed cells in smooth muscle and stromal regions. In contrast, InstanSeg was fast and generally robust, but showed limited ability to resolve certain epithelial subtypes in some regions. µSAM occasionally detected weak nuclear noise as cells, but also performed well in regions with blurred fluorescence signal, indicating good robustness under some challenging imaging conditions. Overall, these results suggest that model choice should depend on the specific imaging characteristics, target cell populations, and downstream biological questions, rather than assuming a single universally optimal segmentation method. (**Table 2, 3, Table S1, 2**).

**Table 3.**
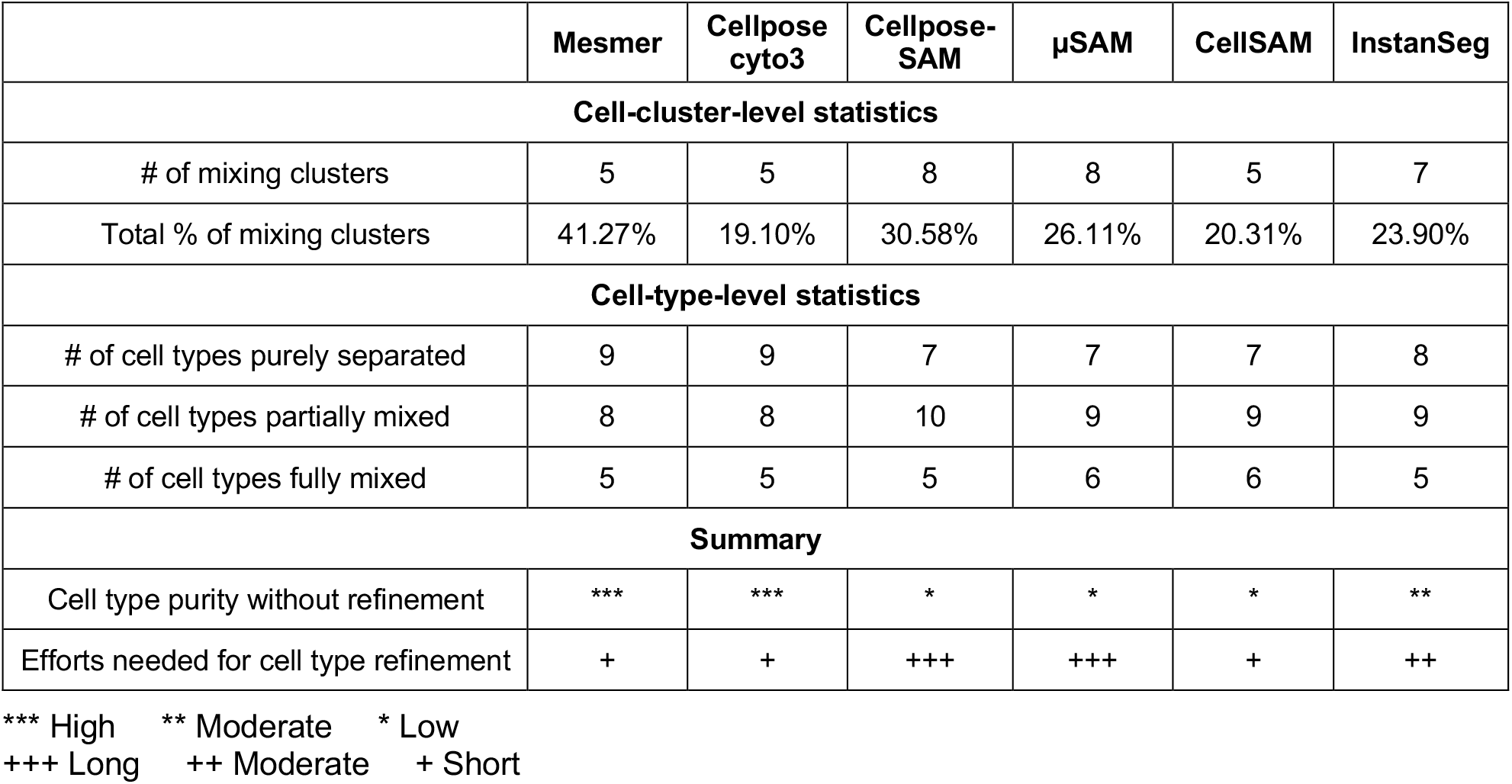
Summary of cell-level segmentation performance across models on multiplexed fluorescence (CODEX) images.

| | Mesmer | Cellpose cyto3 | Cellpose-SAM | μSAM | CellSAM | InstanSeg |
| --- | --- | --- | --- | --- | --- | --- |
| **Cell-cluster-level statistics** |  |  |  |  |  |  |
| # of mixing clusters | 5 | 5 | 8 | 8 | 5 | 7 |
| Total % of mixing clusters | 41.27% | 19.10% | 30.58% | 26.11% | 20.31% | 23.90% |
| **Cell-type-level statistics** |  |  |  |  |  |  |
| # of cell types purely separated | 9 | 9 | 7 | 7 | 7 | 8 |
| # of cell types partially mixed | 8 | 8 | 10 | 9 | 9 | 9 |
| # of cell types fully mixed | 5 | 5 | 5 | 6 | 6 | 5 |
| **Summary** |  |  |  |  |  |  |
| Cell type purity without refinement | *** | *** | * | * | * | ** |
| Efforts needed for cell type refinement | + | + | +++ | +++ | + | ++ |

\*\*\* High \*\* Moderate \* Low +++ Long ++ Moderate + Short

## Discussion

In this study, we systematically evaluated Cellpose cyto3, Cellpose-SAM, µSAM, and CellSAM across multiple imaging modalities, and further examined how segmentation differences propagate into downstream biological analyses. Our results demonstrate that segmentation performance is context-dependent. In live-cell phase imaging data, Cellpose-SAM showed superior performance in delineating cell boundaries. In fluorescence cell culture data, all SAM-based models achieved performance comparable to human annotations and maintained robustness under low signal-to-noise conditions, whereas Cellpose cyto3 was more sensitive to variations in imaging quality and frequently failed to capture accurate cell morphology.

Beyond mask-level evaluation, a central goal of this work was to determine whether differences in segmentation meaningfully affect downstream biological interpretation. By applying each segmentation model to multiplexed fluorescence CODEX datasets and performing clustering and cell type annotation based on extracted marker intensities, we observed that segmentation variability leads to measurable differences in cell type assignments, with all approaches introducing some degree of cell type mixing. Small variations in boundary delineation led to measurable shifts in MUC2 and SOX9 intensities, which in turn altered the separation of goblet and TA cell populations and resulted in markedly different cell type proportions across models. Notably, the persistence of mixed clusters across all segmentation models, despite differences in segmentation strategies, suggests no algorithm is perfect and also that signal contamination between adjacent cells is not solely attributable to segmentation errors (e.g., image registration issues, blurriness of markers). Together, these results demonstrate that segmentation accuracy and generating high quality underlying data are needed for accurate marker quantification and cell type classification in multiplexed spatial proteomics data.

Given these findings, choosing a segmentation model for CODEX or similar multiplexed data requires weighing cluster purity, biological resolution, and practical efficiency against the specific tissue and cell populations of interest. For example, while CellSAM offers one of the best cluster purities, it requires substantially longer computation times. Mesmer represents a reasonable trade-off that although it produced the highest proportion of mixed cells, the total number of mixed cluster types was comparable to CellSAM, meaning the downstream manual refinement required may be similar in practice by an end-user but with less computational time. InstanSeg can be used for large-scale analyses where computational efficiency is critical, though it showed limited ability to resolve certain epithelial subtypes in some regions. µSAM demonstrated the best capability of dealing with lower quality images such as blurred fluorescence signal though it occasionally detects weak nuclear noise as false positive cells. Ultimately, no single model is universally optimal, and model choice should be guided by the specific imaging characteristics, target cell populations, and downstream biological questions (**Table 2, 3**).

These observations also point to directions for improving pipeline robustness beyond model selection. Incorporating signal compensation or spillover correction methods may help mitigate contamination from neighboring cells^20^, and should be considered alongside careful segmentation choice. Normalization strategy and clustering algorithm^21^ could further impact segmentation-induced differences and could be layered into studies in the future. Moreover, the growing availability of automated cell type annotation tools^22–24^ could enable these additional levels of comparison to be completed more rapidly especially for comparing at the biological interpretation level.

Taken together, our study highlights that while modern segmentation models, particularly SAM-based approaches, have improved segmentation performance across diverse imaging modalities, there are still improvements that could be made. Intrinsic imaging artifacts and biological spatial constraints continue to limit single-cell measurement accuracy in ways that are not fully resolved by current methods. We therefore recommend treating segmentation selection as an iterative, empirically validated decision rather than a fixed preprocessing default. To support this, we provide ready-to-use notebooks enabling direct comparison of segmentation models and downstream analytical outcomes on custom datasets, allowing users to identify the approach best suited to their specific applications.

## Methods

### Imaging datasets acquisition

Phase contrast images of the human lung carcinoma cell line A549 were acquired from in-house experiments^25^. Cells were seeded at a density of 7,200 cells per well in a 96-well plate one day prior to imaging to ensure appropriate cell confluency for segmentation evaluation. Images were captured using a multi-camera array microscope^26^ with a 10x/0.3NA objective lens, with a spatial resolution of 0.5 µm per pixel. Phase contrast images of mouse melanoma B16-F10 cells were acquired using the same experimental procedures and imaging settings. The cell culture was maintained within a stage-top incubator (Okolab), and imaged continuously every 10 minutes for 24 hours.

For fluorescence imaging of human lung carcinoma A549 cells, cells were seeded at a density of 7,200 cells per well in a 96-well plate one day prior to fixation and imaging. On the following day, cells were fixed with 4% paraformaldehyde (PFA). After fixation, cells were stained with APC-conjugated anti-CD81 antibody (BioLegend, cat# 349509) for 1 hour in the dark. Subsequently, nuclei were stained with DAPI at a concentration of 16.7 µg/mL (BioLegend, cat# 422801). Fluorescence images were acquired using the Squid microscope^27^ (Cephla Inc.) with a 20x/0.8NA objective lens, corresponding to a spatial resolution of 0.376 µm per pixel. Images were captured using 405 nm excitation for DAPI and 638 nm excitation for APC-conjugated CD81.

Multiplexed fluorescence images generated using CODEX were downloaded from the Human Biomolecular Atlas Program (HuBMAP) portal (https://portal.hubmapconsortium.org). Each pixels represents 0.377 µm in the raw images. Eight tissue regions from the human small and large intestine were analyzed. The corresponding HuBMAP dataset identifiers are HuBMAP ID HBM946.GRVG.379, HBM889.KDGM.632, HBM776.SDCW.837, HBM784.TKZX.992, HBM538.PGFT.538, HBM433.ZLWP.627, HBM842.LQDP.877, HBM657.JVPV.825.

Segmentation masks generated by the HuBMAP processing pipeline using Cytokit were downloaded and used as reference masks for comparison. Cell type annotations based on the Cytokit segmentation outputs were generated by experts within the HuBMAP consortium through manual review and iterative refinement.

### Segmentation

Cell segmentation was performed using six models: Cellpose cyto3, Cellpose-SAM, CellSAM, µSAM, Mesmer, and InstanSeg. Each model was installed and executed in an independent software environment to avoid version conflicts between dependencies.

Cellpose cyto3 and Cellpose-SAM segmentation were performed following the official tutorial provided in the Cellpose GitHub repository (https://github.com/MouseLand/cellpose). CellSAM segmentation was implemented following the instructions provided in the CellSAM repository (https://github.com/vanvalenlab/cellSAM). µSAM segmentation was performed using the AutomaticPromptGenerator class according to the µSAM repository (https://github.com/computational-cell-analytics/micro-sam). Mesmer segmentation was executed through the SPACEc Python package (https://github.com/yuqiyuqitan/SPACEc). InstanSeg segmentation was performed following the official implementation provided in the InstanSeg repository (https://github.com/instanseg/instanseg).

For different imaging modalities, images were preprocessed and formatted to match the input requirements of each segmentation model as described below.

### Phase contrast image segmentation

Phase contrast images were cropped from their original 3072 × 3072 pixels down to 9 individual images of size 1024 × 1024 pixels before segmentation. Each image was provided to the segmentation models as a single-channel two-dimensional image with shape (Height, Width).

### Fluorescence cell culture image segmentation

Fluorescence cell culture images were used at their original resolution of 2084 × 2084 pixels. Images were formatted as three-channel inputs representing membrane signal, nuclear signal, and an empty channel, resulting in image tensors with shape (Height, Width, 3). The order of channels was adjusted as required by each segmentation model.

### CODEX image segmentation

For CODEX datasets, Cytokeratin and CD45 channels were combined to generate a composite membrane channel, while the Hoechst channel was used as the nuclear signal. These channels were formatted into a three-channel input image consisting of membrane signal, nuclear signal, and a blank channel with shape (Height, Width, 3). Channel ordering was adjusted as required by each segmentation model prior to segmentation.

Model-specific parameter settings are provided in **Supplemental Table 3**.

### Fine tuning of Cellpose-SAM

Fine-tuning of Cellpose-SAM for phase contrast images of A549 cells was performed following the protocol provided in the Cellpose repository. The training dataset consisted of ground-truth segmentation masks manually annotated by human experts from six images (1024×1024 pixels). Model training was performed using the default training parameters provided by Cellpose-SAM. For validation, three independent images with corresponding expert-annotated masks that were not used during training, were employed to evaluate performance improvements after fine-tuning.

### Metrics

The performance of cell segmentation models was quantitatively assessed using standard evaluation metrics, including Intersection over Union (IoU), Average Precision (AP), Segmentation Accuracy (SEG), and F1 score. The definitions are as follows:

Intersection over Union (IoU):

IoU is defined as the ratio of the area of overlap to the area of union between a predicted mask and the corresponding ground truth mask:

Let *G* = {*g*_1_, *g*_2_, …, *g*_*n*_} be the set of ground truth masks and *P* = {*p*_1_, *p*_2_, …, *p*_*m*_} the set of predicted masks. For a given ground truth mask *g*_*i*_, define the IoU with each predicted mask *p*_*j*_ as:

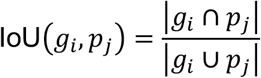

Average Precision (AP):

Average Precision is adapted from Cellpose evaluation metrics^6^. It measures the proportion of correctly predicted masks relative to all predictions and ground truth instances:

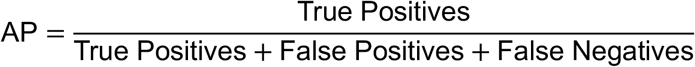

AP is computed across varying IoU thresholds. Unless otherwise specified, an IoU threshold of 0.5 is used, following the general standard established by the PASCAL VOC benchmark^15^. Optimal one-to-one matching between predicted and ground truth masks is performed using the linear sum assignment algorithm. A matched pair is counted as a true positive if their IoU is greater than or equal to the threshold. Unmatched predictions are considered false positives, and unmatched ground truth masks are counted as false negatives.

Segmentation Accuracy (SEG)^16^:

The SEG score quantifies segmentation quality by averaging the Intersection over Union (IoU) values of ground truth–prediction mask pairs.

Let *G* = {*g*_1_, *g*_2_, …, *g*_*n*_} be the set of ground truth masks and *P* = {*p*_1_, *p*_2_, …, *p*_*m*_} the set of predicted masks. The matching between ground truth and predicted masks is computed using the linear sum assignment algorithm. For all each matched pair (*g*_*i*_, *p*_*j*_) ∈ *M*, compute the IoU as defined above. The SEG score is then defined as the mean IoU across all matched pairs, normalized by the number of ground truth instances:

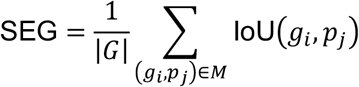

Where:

|*G*| is the number of ground truth masks, *M* is the set of matched ground truth–prediction pairs (true positives), IoU(*g*_*i*_, *p*_*j*_) is the IoU between predicted mask *p* and ground truth mask *g*.

F1 Score:

The F1 score provides a harmonic mean of precision and recall:

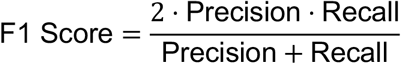

Precision and recall are defined as:

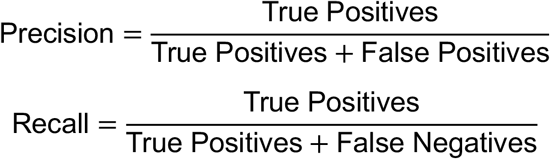

A predicted mask is considered a true positive if it matches a ground truth mask with an IoU greater than or equal to a specified threshold (e.g., 0.5). Unless otherwise specified, an IoU threshold of 0.5 is used. Unmatched predicted masks are counted as false positives, and unmatched ground truth masks are counted as false negatives.

### Mask matching for CODEX datasets

To evaluate the concordance between segmentation masks generated by different models on the CODEX datasets, we performed pairwise mask matching between segmentation results from each model pair.

For each mask from model A, we identified candidate masks from model B whose centroids were located within a 10-pixel radius of the centroid of the mask from model A. Among these candidates, the nearest mask from model B was selected. If the corresponding mask from model B also identified the mask from model A as its nearest neighbor, the two masks were defined as a mutual nearest neighbor pair and considered a matched mask.

### Processing and annotation of CODEX datasets

Segmentation outputs from all models were processed using the same analysis pipeline. For each segmentation mask, the mean intensity of every protein marker channel was calculated and used to generate a feature matrix for each cell. The centroid coordinates (x, y) of each mask were also recorded.

Feature matrices from all eight tissue regions were merged into a single dataset. Marker intensities were then z-normalized for each marker across all cells, except for metadata columns and the two nuclear markers DRAQ5 and Hoechst1.

Three noise filtering steps were applied to remove low-quality cell masks. First, cells with extreme overall signal values were filtered out. Specifically, cells were removed if the sum of z-scores across all markers exceeded 20 and if more than 20 markers had a z-score greater than 1. Second, cells with very weak nuclear signals were excluded using the criteria DRAQ5 < 80 and Hoechst1 < 800. Third, masks with abnormal sizes were removed by retaining only cells with areas between 40 and 2000 pixels.

The resulting feature matrix was clustered using unsupervised Leiden clustering based on the 45-marker panel. Clustering was performed using the rapids-singlecell Python package (https://github.com/scverse/rapids-singlecell). For each segmentation model, the resulting clusters were manually annotated as either pure clusters or mixed clusters. This annotation was performed by visually overlaying cells belonging to each cluster onto the raw CODEX images in the Napari software and examining marker expression patterns and cell type purity.

To further assess the impact of segmentation on resolving epithelial subtypes, we performed a mock subclustering analysis on mixed epithelial clusters within a representative epithelial region. Specifically, clusters annotated as “goblet/TA/other epithelial” and “TA/other epithelial” were re-clustered into three groups using unsupervised k-means clustering (k = 3). Subclusters with high SOX9 expression were labeled as transit-amplifying (TA) cells, those with high MUC2 expression were labeled as goblet cells, and the remaining cells were labeled as other epithelial cell types.

## Data availability

CODEX raw images and processed datasets are available through the Human Biomolecular Atlas Program (HuBMAP) portal (https://portal.hubmapconsortium.org). Human expert-annotated CODEX datasets, segmentation outputs from all models, and raw phase contrast and fluorescence cell culture images are available from the Duke Research Repository^28^.

## Code availability

All code used for figure generation, as well as notebooks for comparing segmentation models on user datasets, is available in our GitHub repository (https://github.com/HickeyLab/Segmentation_Comparison).

## Author contributions

Y.M. and J.L. acquired the imaging data. Y.M., N.S., J.L., K.P., K.A. performed segmentation. Y.M., N.S., J.L., K.A., Y.L., Y.Z., K.VB., J.J., J.vdK., B.YX.N., A.A. and R.R. performed data analysis. Y.M., J.L. and J.W.H wrote the manuscript. R.H. and J.W.H. supervised this study.

## Acknowledgements

This work was supported by the US National Institutes of Health (grant nos. 3U54AG07593, 3OT2OD033759-01S4, 3OT2OD033759-01S5, 1U01-AI186999-01, 1U54-AI191253-01); the National Science Foundation CAREER Award (grant no. 2440733); the V Foundation (V2025-019); the Human Frontier Science Program Early Career Grant (RGEC26/2025-); and start-up support from Duke University Department of Biomedical Engineering for JWH.

## References

1. Seal, S. et al. Cell Painting: a decade of discovery and innovation in cellular imaging. Nat Methods 22, 254–268 (2025).

2. Park, Y., Depeursinge, C. & Popescu, G. Quantitative phase imaging in biomedicine. Nature Photon 12, 578–589 (2018).

3. Bressan, D., Battistoni, G. & Hannon, G. J. The dawn of spatial omics. Science 381, eabq4964 (2023).

4. Liu, Y., Dai, Y. & Wang, L. Spatial omics at the forefront: emerging technologies, analytical innovations, and clinical applications. Cancer Cell 44, 24–49 (2026).

5. Black, S. et al. CODEX multiplexed tissue imaging with DNA-conjugated antibodies. Nat Protoc 16, 3802–3835 (2021).

6. Stringer, C., Wang, T., Michaelos, M. & Pachitariu, M. Cellpose: a generalist algorithm for cellular segmentation. Nat Methods 18, 100–106 (2021).

7. Pachitariu, M. & Stringer, C. Cellpose 2.0: how to train your own model. Nat Methods 19, 1634–1641 (2022).

8. Stringer, C. & Pachitariu, M. Cellpose3: one-click image restoration for improved cellular segmentation. Nat Methods 22, 592–599 (2025).

9. Pachitariu, M., Rariden, M. & Stringer, C. Cellpose-SAM: superhuman generalization for cellular segmentation. Preprint at 10.1101/2025.04.28.651001 (2025).

10. Greenwald, N. F. et al. Whole-cell segmentation of tissue images with human-level performance using large-scale data annotation and deep learning. Nat Biotechnol 40, 555–565 (2022).

11. Goldsborough, T. et al. InstanSeg: an embedding-based instance segmentation algorithm optimized for accurate, efficient and portable cell segmentation. Preprint at 10.48550/ARXIV.2408.15954 (2024).

12. Kirillov, A. et al. Segment Anything. Preprint at 10.48550/ARXIV.2304.02643 (2023).

13. Archit, A. et al. Segment Anything for Microscopy. Nat Methods 22, 579–591 (2025).

14. Marks, M. et al. CellSAM: a foundation model for cell segmentation. Nat Methods 22, 2585–2593 (2025).

15. Everingham, M., Van Gool, L., Williams, C. K. I., Winn, J. & Zisserman, A. The Pascal Visual Object Classes (VOC) Challenge. Int J Comput Vis 88, 303–338 (2010).

16. Maška, M. et al. The Cell Tracking Challenge: 10 years of objective benchmarking. Nat Methods 20, 1010–1020 (2023).

17. Hickey, J. W. et al. Organization of the human intestine at single-cell resolution. Nature 619, 572–584 (2023).

18. Czech, E., Aksoy, B. A., Aksoy, P. & Hammerbacher, J. Cytokit: a single-cell analysis toolkit for high dimensional fluorescent microscopy imaging. BMC Bioinformatics 20, 448 (2019).

19. Tan, Y. et al. SPACEc: a streamlined, interactive Python workflow for multiplexed image processing and analysis. Nat Commun 16, 10652 (2025).

20. Bai, Y. et al. Adjacent Cell Marker Lateral Spillover Compensation and Reinforcement for Multiplexed Images. Front. Immunol. 12, 652631 (2021).

21. Hickey, J. W., Tan, Y., Nolan, G. P. & Goltsev, Y. Strategies for Accurate Cell Type Identification in CODEX Multiplexed Imaging Data. Front. Immunol. 12, 727626 (2021).

22. Rumberger, J. L. et al. Automated classification of cellular expression in multiplexed imaging data with Nimbus. Nat Methods 22, 2161–2170 (2025).

23. Wang, X. (Julie) et al. Generalized cell phenotyping for spatial proteomics with language-informed vision models. Preprint at 10.1101/2024.11.02.621624 (2024).

24. Sun, H. et al. Flexible and robust cell-type annotation for highly multiplexed tissue images. Cell Systems 16, 101374 (2025).

25. Lerner, J. F. et al. Extremely high-throughput quantitative phase imaging of live-cells with a multi-camera array microscope. in Quantitative Phase Imaging XII (eds Park, Y. & Liu, Y.) 3 (SPIE, lSan Francisco, United States, 2026). doi:10.1117/12.3077986.

26. Kim, K. et al. Rapid 3D imaging at cellular resolution for digital cytopathology with a multi-camera array scanner (MCAS). npj Imaging 2, 39 (2024).

27. Hickey, J. W. et al. Fluid-Squid: DIY Multiplexed Imaging of Cells and Tissues. Preprint at 10.1101/2025.10.09.680291 (2025).

28. Miao, Y. & Hickey, J. Data from: Foundation cell segmentation models performance on live microscopy and spatial-omics data. 5.6 GB Duke Research Data Repository 10.7924/R4R505 (2026).

