## Supplemental Information for "Foundation cell segmentation models performance on live microscopy and spatial-omics data"

**This file includes:**

Supplemental Table 1-3

Supplemental Figure 1-5

Supplemental File 1-7

| **Category** | **Cluster name** | **Mesmer** | **Cellpose cyto3** | **Cellpose-SAM** | **µSAM** | **CellSAM** | **InstanSeg** |
| --- | --- | --- | --- | --- | --- | --- | --- |
| Pure | Nerve | V | V | V | V | V | V |
| Pure | Neuroendocrine | V | V | V | V | V | V |
| Pure | Neutrophil | V | V | V | V | V | V |
| Pure | Paneth | V | V | V | V | V | V |
| Pure | Plasma | V | V | V | V | V | V |
| Pure | Smooth muscle | V | V | V | V | V | V |
| Pure | Stromal | V | V | V | V | V | V |
| Pure | Endothelial | V | V | V | V | V | V |
| Pure | Epithelial | V | V | V | V | V | V |
| Pure | CD66+ epithelial | V | V | V | V | V | V |
| Pure | B | V | V | V | V | V | V |
| Pure | CD7+ immune | V | V | V | V | V | V |
| Pure | CD8+ T | V | V | V | V | V | V |
| Pure | M2 macrophage | V | V | V | V | V | V |
| Pure | M1 macrophage | V |  | V | V | V | V |
| Pure | CD4+ T |  | V |  | V | V |  |
| Pure | Cycling TA |  |  | V |  | V |  |
| Pure | DC |  |  |  |  | V | V |
| Pure | ICC | V |  |  |  |  |  |
| Pure | Goblet |  |  |  |  |  |  |
| Pure | TA |  |  |  |  |  |  |
| Pure | NK |  |  |  |  |  |  |
| Mixed | Goblet/TA/Other epithelial | V | V | V | V | V | V |
| Mixed | Immune mixed | V | V | V | V | V | V |
| Mixed | B/CD4+ T mixed | V | V | V | V | V | V |
| Mixed | Stromal/immune mixed |  |  | V | V | V | V |
| Mixed | Goblet/TA/CD66+/Other epithelial |  | V | V | V |  |  |
| Mixed | DC/Macrophage/  CD4+ T mixed |  | V | V | V |  |  |
| Mixed | CD4+ T/CD8+ T mixed | V |  | V |  |  | V |
| Mixed | TA/Other epithelial |  |  | V |  |  | V |
| Mixed | Imaging artifacts/Stromal | V |  |  | V |  |  |
| Mixed | Nerve/immune mixed |  |  |  | V | V |  |
| Mixed | Stromal/epithelial mixed |  |  |  |  |  | V |

A check mark (V) indicates the presence of the corresponding cluster in each model.

**Table S1.** Summary of Leiden cluster occurrence across segmentation models on multiplexed fluorescence (CODEX) images.

| **Cell type** | **Mesmer** | **Cellpose cyto3** | **Cellpose-SAM** | **µSAM** | **CellSAM** | **InstanSeg** |
| --- | --- | --- | --- | --- | --- | --- |
| **Epithelial** | | | | | | |
| Epithelial (MUC1+, ITLN1+, non-specialized) | Partially mixed | Partially mixed | Partially mixed | Partially mixed | Partially mixed | Partially mixed |
| Goblet | Mixed | Mixed | Mixed | Mixed | Mixed | Mixed |
| TA | Mixed | Mixed | Mixed | Mixed | Mixed | Mixed |
| Cycling TA | Mixed | Mixed | Partially mixed | Mixed | Mixed | Mixed |
| CD66+ epithelial | Pure | Partially mixed | Partially mixed | Partially mixed | Partially mixed | Pure |
| Paneth | Pure | Pure | Pure | Pure | Pure | Pure |
| Neuroendocrine | Pure | Pure | Pure | Pure | Pure | Pure |
| **Immune** | | | | | | |
| CD8+ T | Partially mixed | Pure | Partially mixed | Pure | Pure | Partially mixed |
| CD4+ T | Partially mixed | Partially mixed | Partially mixed | Partially mixed | Partially mixed | Partially mixed |
| B | Partially mixed | Partially mixed | Partially mixed | Partially mixed | Partially mixed | Partially mixed |
| Plasma | Partially mixed | Partially mixed | Partially mixed | Partially mixed | Partially mixed | Partially mixed |
| Dendric cells | Mixed | Partially mixed | Mixed | Mixed | Mixed | Partially mixed |
| M1 macrophage | Partially mixed | Partially mixed | Partially mixed | Partially mixed | Partially mixed | Partially mixed |
| M2 macrophage | Partially mixed | Partially mixed | Partially mixed | Partially mixed | Partially mixed | Partially mixed |
| Neutrophil | Pure | Pure | Pure | Pure | Pure | Pure |
| CD7+ immune | Pure | Pure | Pure | Pure | Pure | Pure |
| NK | Mixed | Mixed | Mixed | Mixed | Mixed | Mixed |
| **Stromal** | | | | | | |
| Stromal | Partially mixed | Pure | Partially mixed | Partially mixed | Partially mixed | Partially mixed |
| Smooth muscle | Pure | Pure | Pure | Pure | Pure | Pure |
| Endothelial | Pure | Pure | Pure | Pure | Pure | Pure |
| Nerve | Pure | Pure | Pure | Partially mixed | Partially mixed | Pure |
| ICC | Pure | Mixed | Mixed | Mixed | Mixed | Mixed |

**Pure**: This cell type forms a distinct cluster without mixing with other cell types.

**Partially mixed**: This cell type forms a distinct cluster but is also present in mixed clusters.

**Mixed**: This cell type appears only in mixed clusters with other cell types.

**Table S2.** Summary of cell type mixing across segmentation models on multiplexed fluorescence (CODEX) images.

|  | **Modality** | **model version/name** | **Segmentation parameters**  **(others are default)** | **Fluorescence channel information** |
| --- | --- | --- | --- | --- |
| **Cellpose cyto3** | Phase contrast | cyto3 | channels=[0,0] |  |
|  |  |  | diameter=0 |  |
|  | Fluorescence cell culture | cyto3 | channels=[1, 3] | Rearrange channel order to be [membrane, blank, nucleus] |
|  |  |  | diameter=0 |  |
|  |  |  | flow_threshold=0.7 |  |
|  | CODEX | cyto3 | channels=[1, 3] | Rearrange channel order to be [membrane, blank, nucleus] |
|  |  |  | diameter=0 |  |
|  |  |  | flow_threshold=0.7 |  |
| **Cellpose-SAM** | Phase contrast | cpsam | batch_size=32 |  |
|  |  |  | flow_threshold = 0.7 |  |
|  |  |  | cellprob_threshold = 0.0 |  |
|  |  |  | tile_norm_blocksize = 0 |  |
|  | Fluorescence cell culture | cpsam | batch_size=32 | Rearrange channel order to be [blank, nucleus, membrane] |
|  |  |  | flow_threshold = 0.7 |  |
|  |  |  | cellprob_threshold = 0.0 |  |
|  |  |  | tile_norm_blocksize = 0 |  |
|  | CODEX | cpsam | batch_size=32 | Rearrange channel order to be [blank, nucleus, membrane] |
|  |  |  | flow_threshold = 0.7 |  |
|  |  |  | cellprob_threshold = 0.0 |  |
|  |  |  | tile_norm_blocksize = 0 |  |
| **µSAM** | Phase contrast | vit_l_lm (AutomaticPromptGenerator) | ndim=2 |  |
|  |  |  | tile_shape=(1024, 1024) |  |
|  |  |  | halo=(256, 256) |  |
|  | Fluorescence cell culture | vit_l_lm (AutomaticPromptGenerator) | ndim=2 | Rearrange channel order to be [membrane, nucleus, blank] |
|  | CODEX | vit_l_lm (AutomaticPromptGenerator) | tile_shape=(512, 512) | Rearrange channel order to be [membrane, nucleus, blank] |
|  |  |  | halo=(64, 64) |  |
|  |  |  | optimize_memory=True |  |
| **CellSAM** | Phase contrast | cellsam_base_v1.1 | block_size=256 |  |
|  |  |  | overlap=100 |  |
|  |  |  | iou_depth=50 |  |
|  |  |  | low_contrast_enhancement=True |  |
|  |  |  | use_wsi=True |  |
|  | Fluorescence cell culture | cellsam_base_v1.1 | chunks=256 | Rearrange channel order to be [blank, nucleus, membrane] |
|  |  |  | block_size=1024 |  |
|  |  |  | overlap=300 |  |
|  |  |  | iou_depth=50 |  |
|  |  |  | low_contrast_enhancement=False |  |
|  |  |  | use_wsi=True |  |
|  |  |  | gauge_cell_size=False |  |
|  | CODEX | cellsam_base_v1.1 | block_size=250 | Rearrange channel order to be [blank, nucleus, membrane] |
|  |  |  | overlap=100 |  |
|  |  |  | iou_depth=100 |  |
|  |  |  | low_contrast_enhancement=False |  |
|  |  |  | use_wsi=True |  |
|  |  |  | gauge_cell_size=True |  |
| **Mesmer**  **(SPACEc package)** | CODEX | default | seg_method='mesmer' | Channel_0 is corresponding to Hoechst;  Channel_41 is corresponding to Cytokeratin;  Channel_45 is corresponding to CD45 |
|  |  |  | nuclei_channel="Channel_0" |  |
|  |  |  | membrane_channel_list=["Channel_41", "Channel_48"] |  |
|  |  |  | compartment='whole-cell' |  |
|  |  |  | input_format='Multichannel' |  |
|  |  |  | resize_factor=0.7 |  |
|  |  |  | size_cutoff=0 |  |
| **InstanSeg** | CODEX | fluorescence_nuclei_and_cells version 0.1.0 | channel_id = [2,0] | Rearrange channel order to be [membrane, blank, nucleus] |
|  |  |  | pixel_size=0.5 |  |
|  |  |  | resolve_cell_and_nucleus=True, |  |
|  |  |  | cleanup_fragments = True |  |

**Table S3.** Summary of parameter settings for each segmentation model across phase contrast, fluorescence cell culture, and multiplexed fluorescence (CODEX) imaging modalities.

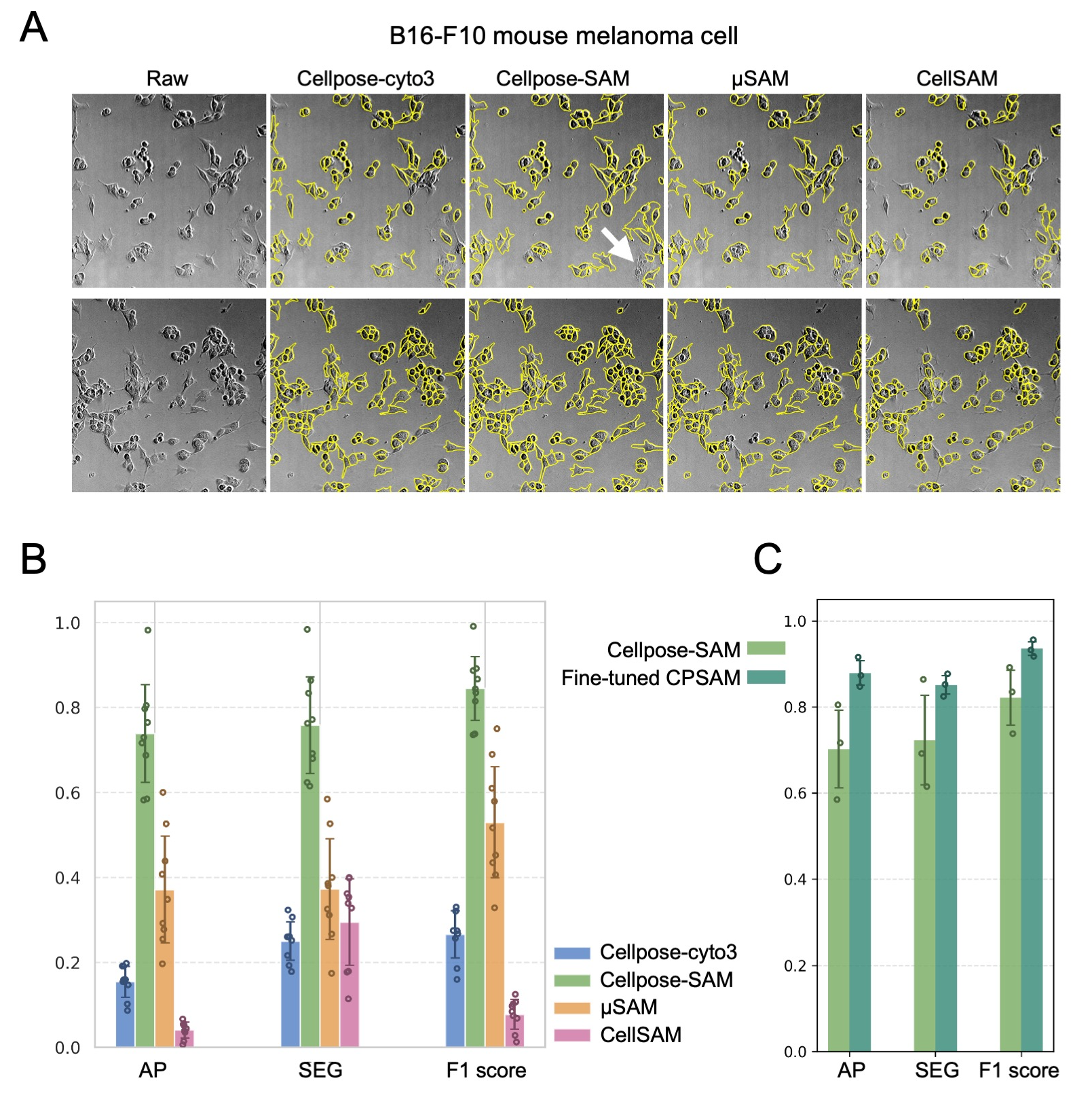

**Supplemental Figure 1.** **Model comparison on phase contrast images.** (A) Representative overlays of segmentation masks for B16-F10 cells on raw images; mask boundaries are shown in yellow, with Cellpose-SAM errors indicated by white arrows.

(B) Performance metrics (AP, SEG, F1 score) on the A549 dataset across models.

(C) Comparison of performance metrics between the Cellpose-SAM base and fine-tuned models on the A549 dataset.

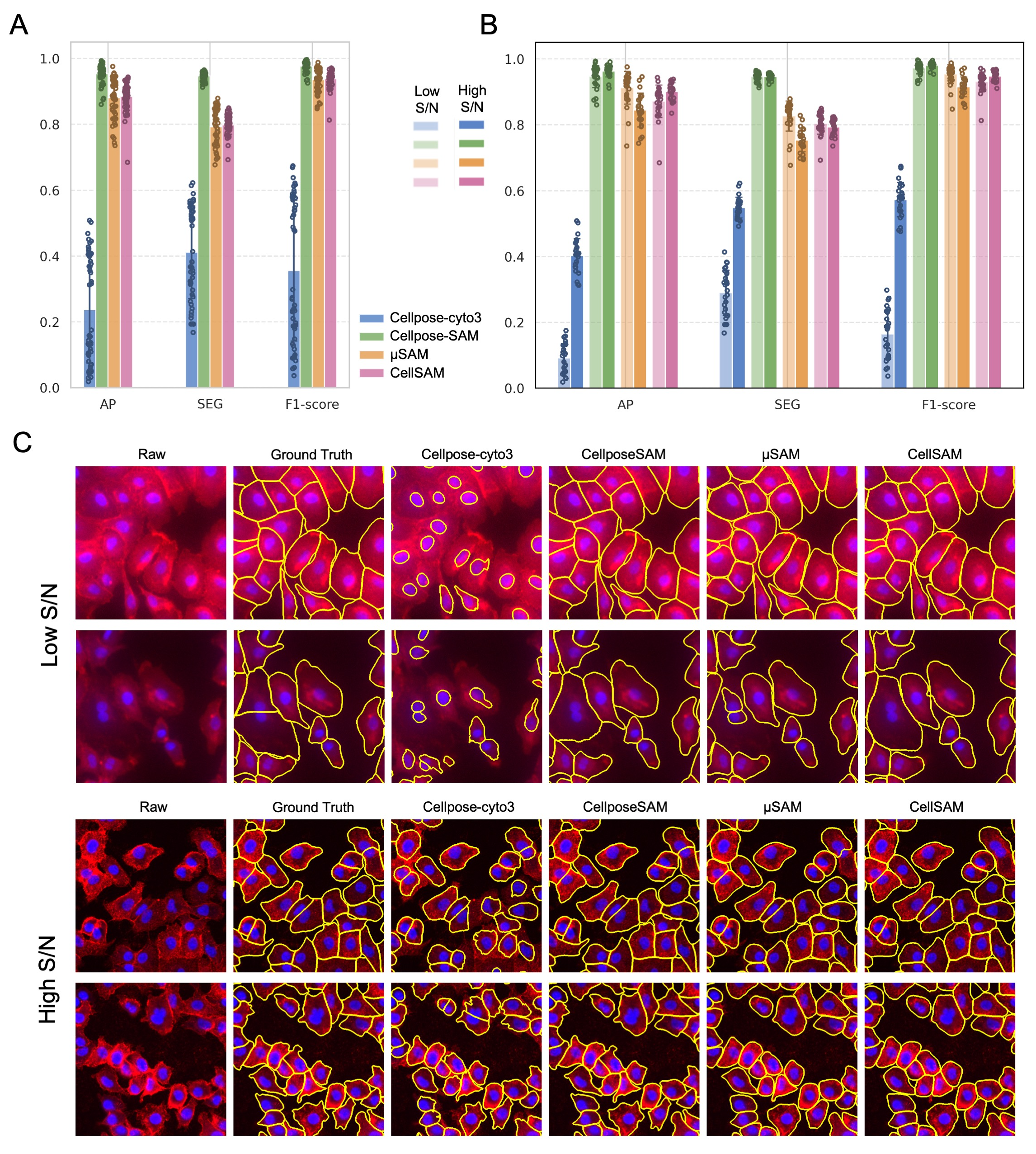

**Supplemental Figure 2.** **Model comparison on fluorescence cell culture images.** (A) Performance metrics (AP, SEG, F1 score) on the A549 dataset across models. (B) Performance metrics grouped by high and low signal-to-noise (S/N) conditions. (C) Representative overlays of segmentation masks under low and high S/N conditions, illustrating differences in Cellpose cyto3 performance; mask boundaries are shown in yellow.

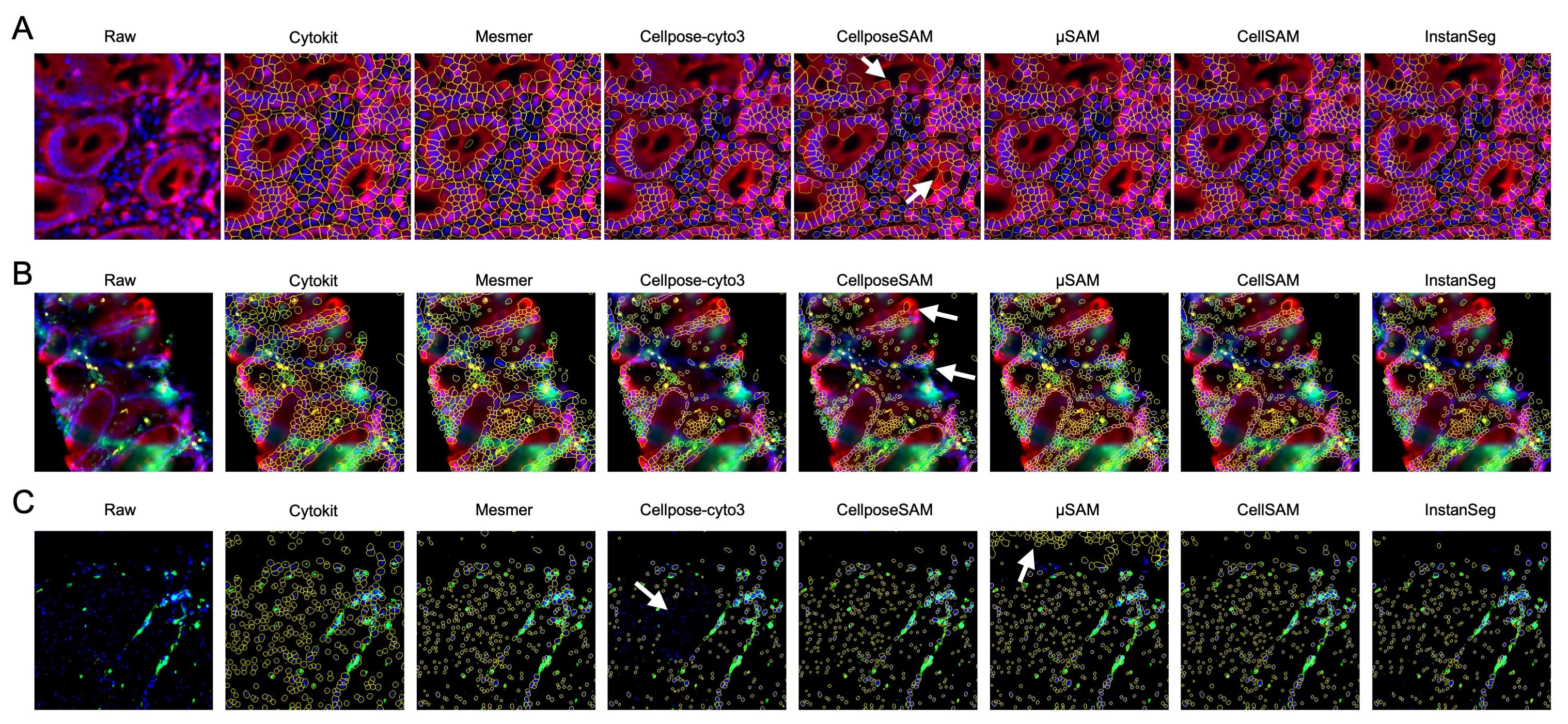

**Supplementary Figure 3. Model comparison on multiplexed fluorescence CODEX images in challenging regions.** (A) Representative overlays of segmentation masks in an epithelial region with weak membrane (cytokeratin) signal. Cellpose-SAM errors are indicated by white arrows. Nucleus is shown in blue, cytokeratin in red; mask boundaries are shown in yellow. (B) Representative overlays in a region with signal blur. Missing masks from Cellpose-SAM are indicated by white arrows. Nucleus is shown in blue, cytokeratin in red, and CD45 in green; mask boundaries are shown in yellow. (C) Representative overlays highlighting model-specific errors, including missed detections by Cellpose cyto3 and over-segmentation by µSAM (white arrows).

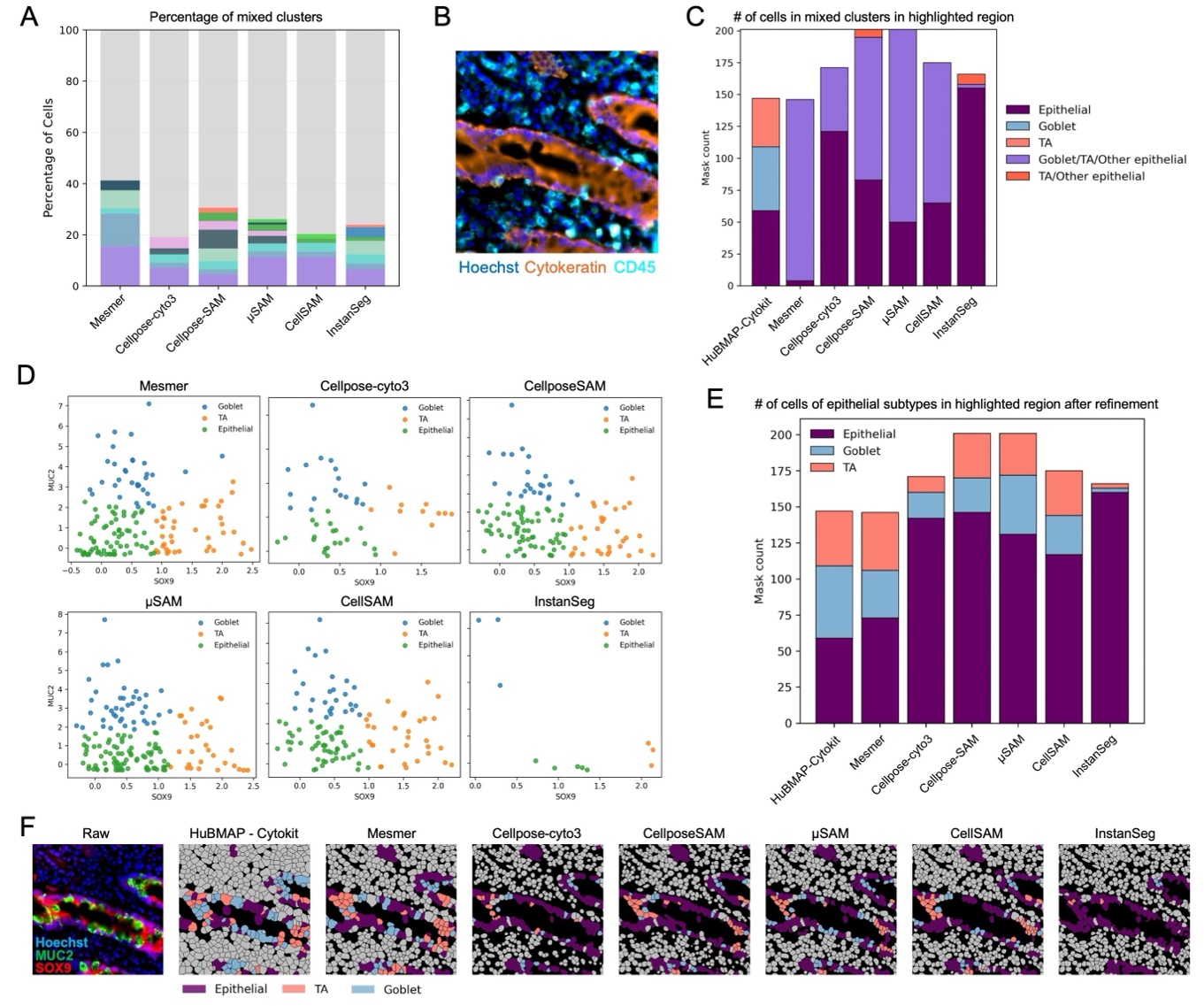

**Supplementary Figure 4. Model comparison on multiplexed fluorescence CODEX images at the level of downstream cell type classification.** (A) Proportion of mixed clusters per model, with mixed clusters stacked to highlight total percentages. (B) Raw image region corresponding to Fig. 5, showing channels used for segmentation. Nucleus is shown in blue and cytokeratin in orange; mask boundaries are shown in cyan. (C) Quantification of epithelial subtypes in the representative region from Fig. 5. (D) k-means subclustering (k = 3) of mixed epithelial clusters using SOX9 and MUC2 expression across models. (E) Quantification of epithelial subtypes after k-means refinement. (F) Segmentation masks mapped to tissue space in the representative region from Fig. 5, colored by cell type after k-means refinement. The raw image (left) shows nucleus in blue, TA marker SOX9 in red, and goblet marker MUC2 in green.

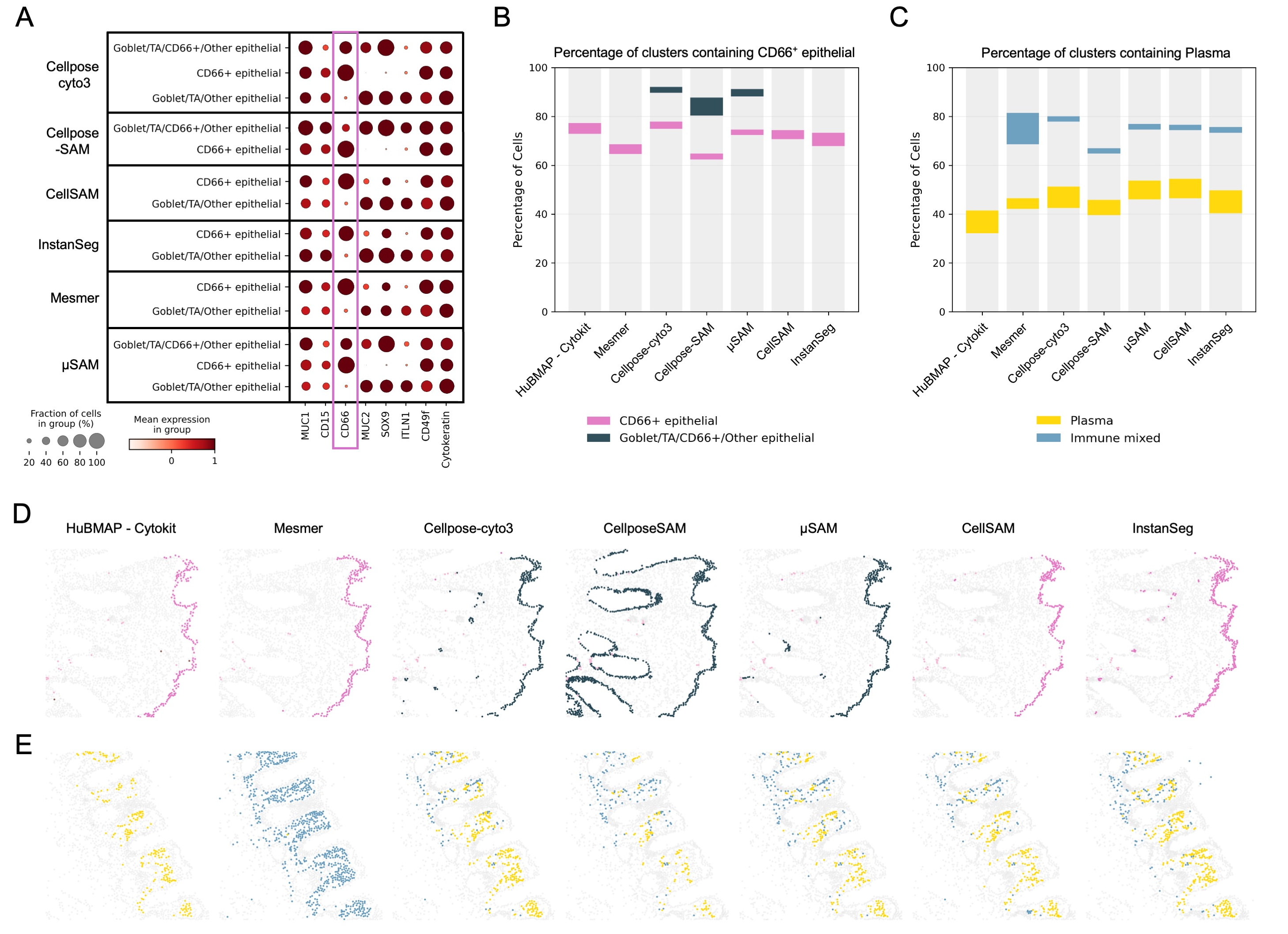

**Supplementary Figure 5. Additional examples of cell type mixing in CODEX images.** (A) Marker expression profiles of epithelial markers in clusters containing CD66+ epithelial cells, highlighting differences in mixing across models. (B) Proportion of CD66+ epithelial clusters and mixed clusters containing CD66+ epithelial cells. (C) Proportion of plasma cell clusters and mixed clusters containing plasma cells (immune mixed clusters). (D) Representative tissue region illustrating CD66+ epithelial clusters and associated mixed clusters, corresponding to (B). (E) Representative tissue region illustrating plasma cell clusters and associated mixed clusters, corresponding to (C).

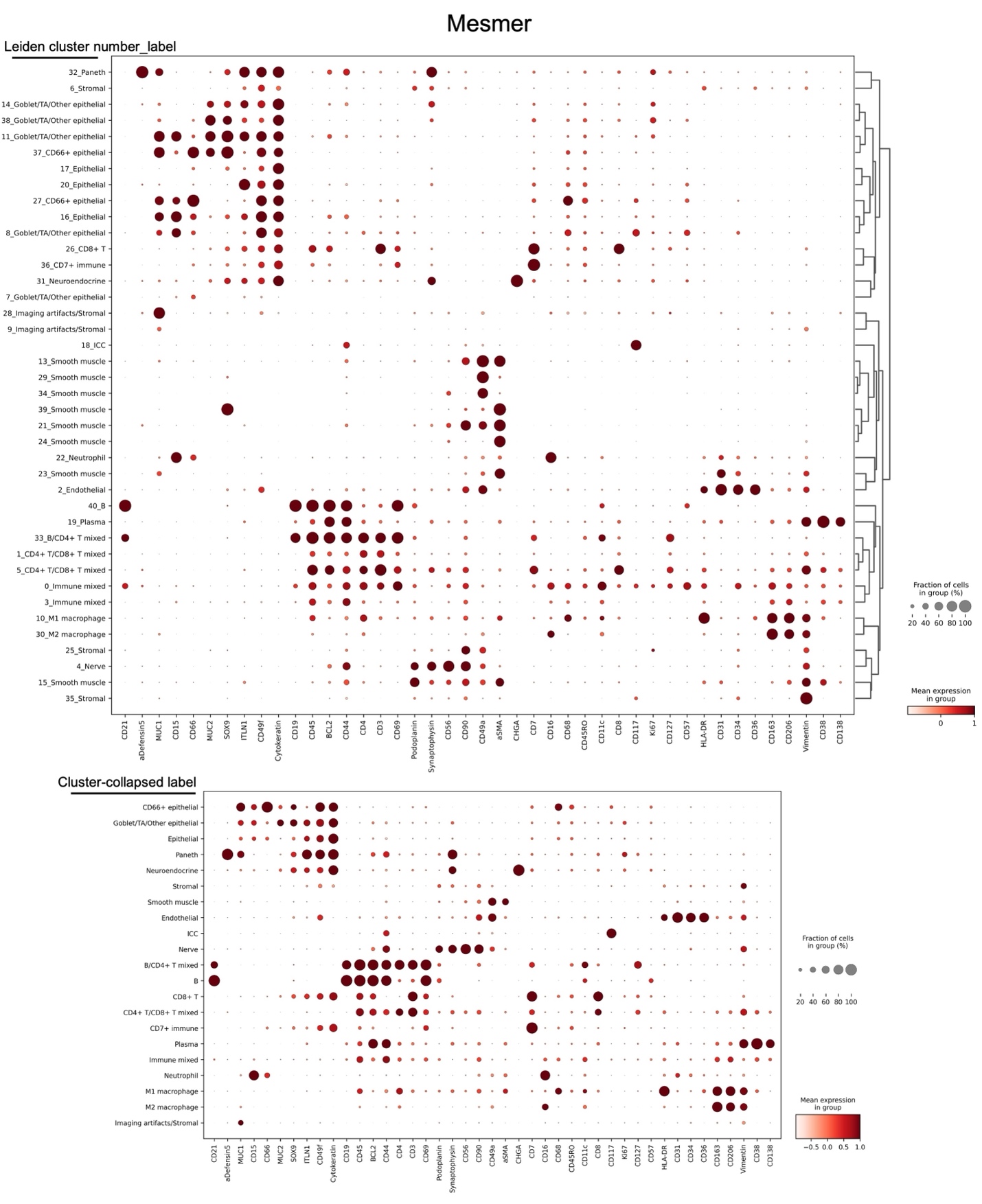

**Supplementary File 1. Leiden cluster labels before and after merging of identical labels for Mesmer segmentation outputs.**

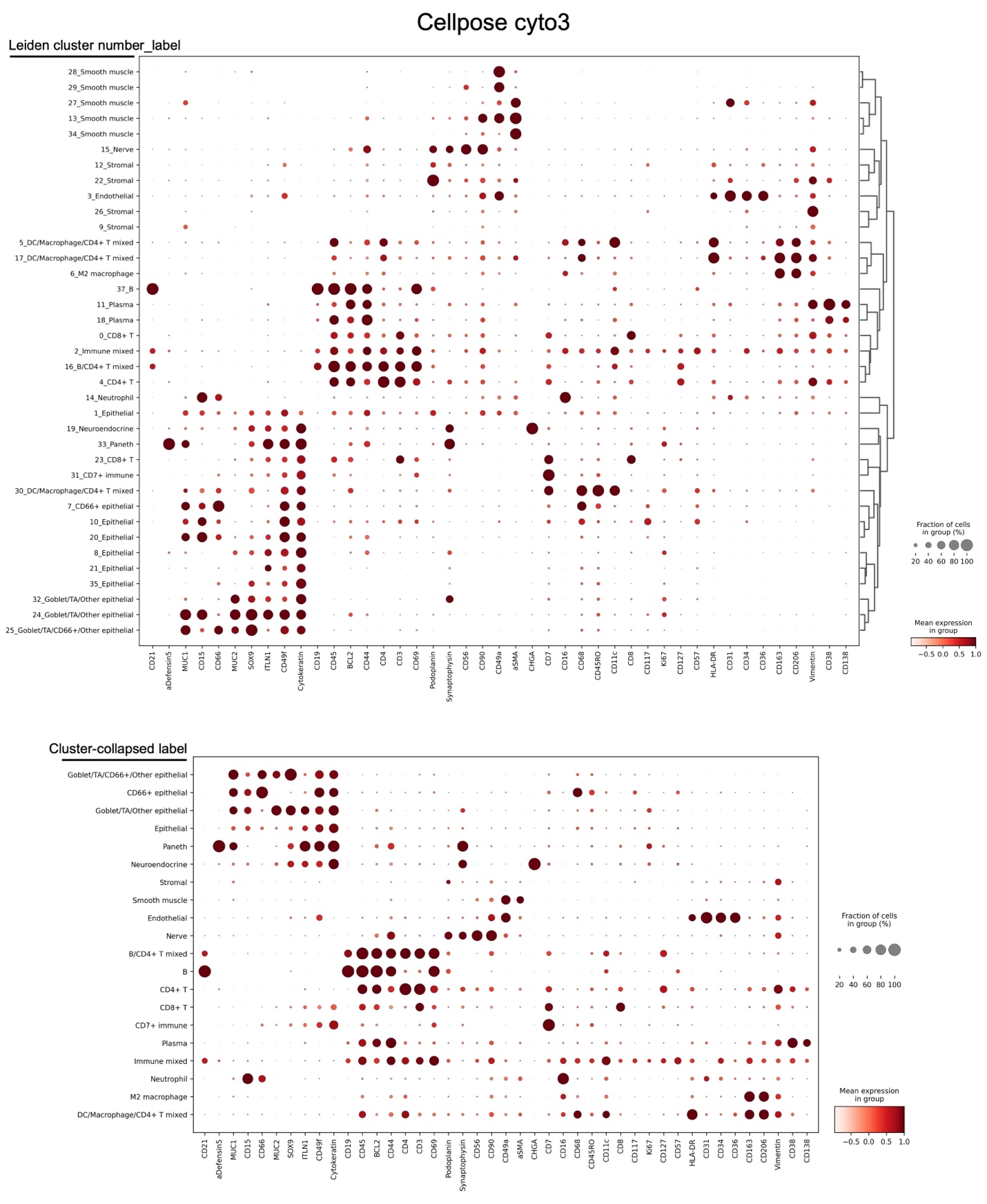
**Supplementary File 2. Leiden cluster labels before and after merging of identical labels for Cellpose cyto3 segmentation outputs.**

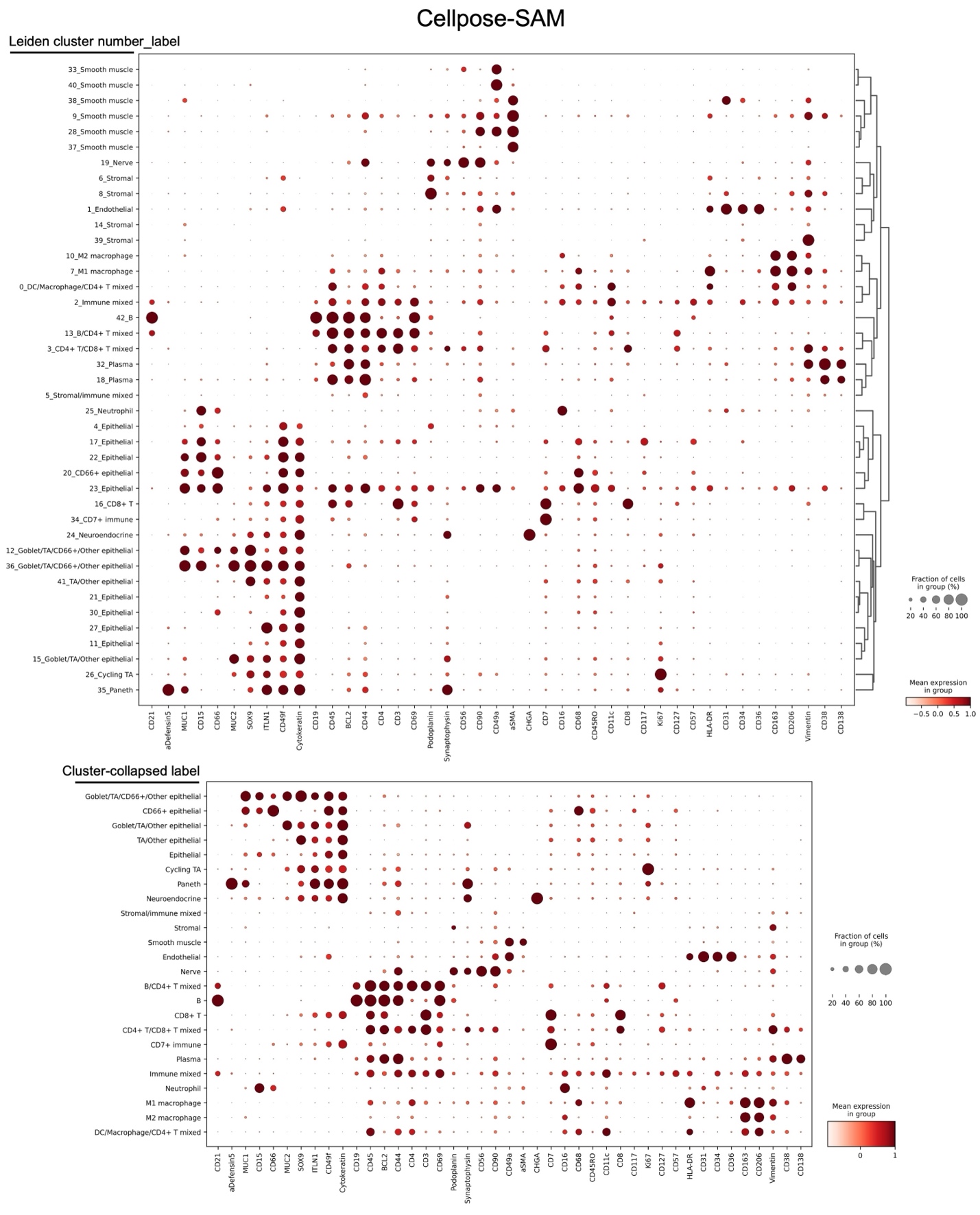

**Supplementary File 3. Leiden cluster labels before and after merging of identical labels for Cellpose-SAM segmentation outputs.**

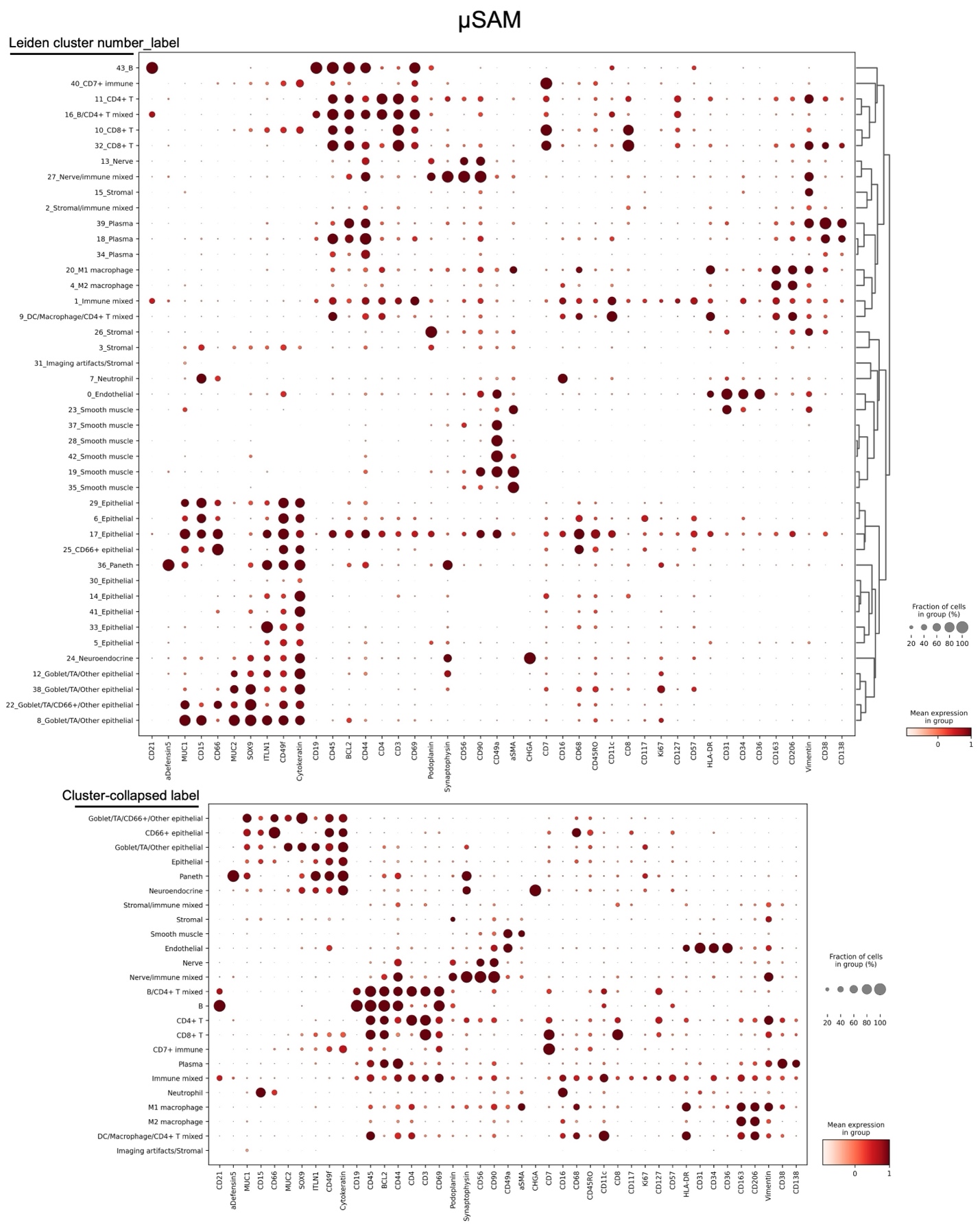

**Supplementary File 4. Leiden cluster labels before and after merging of identical labels for µSAM segmentation outputs.**

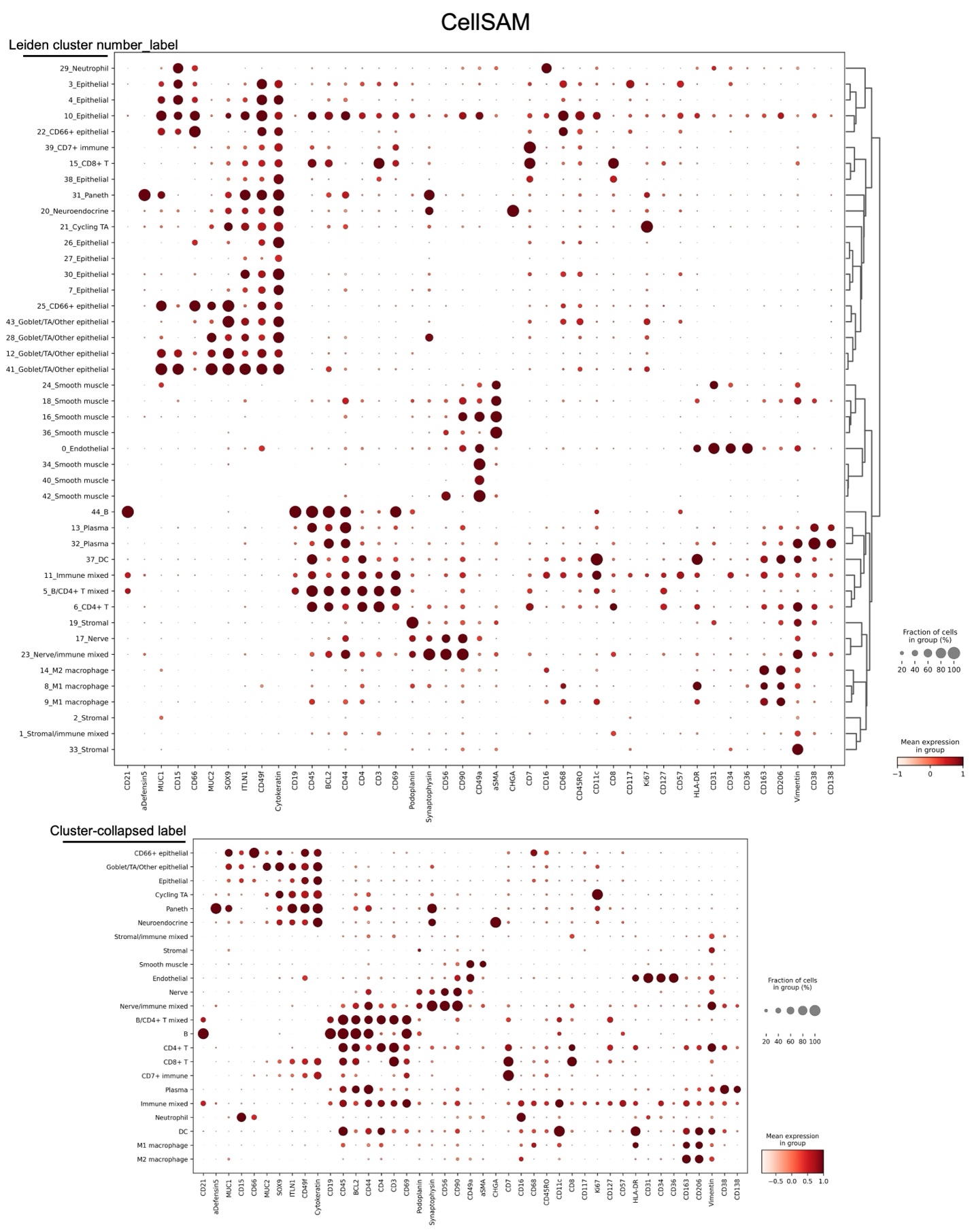

**Supplementary File 5. Leiden cluster labels before and after merging of identical labels for CellSAM segmentation outputs.**

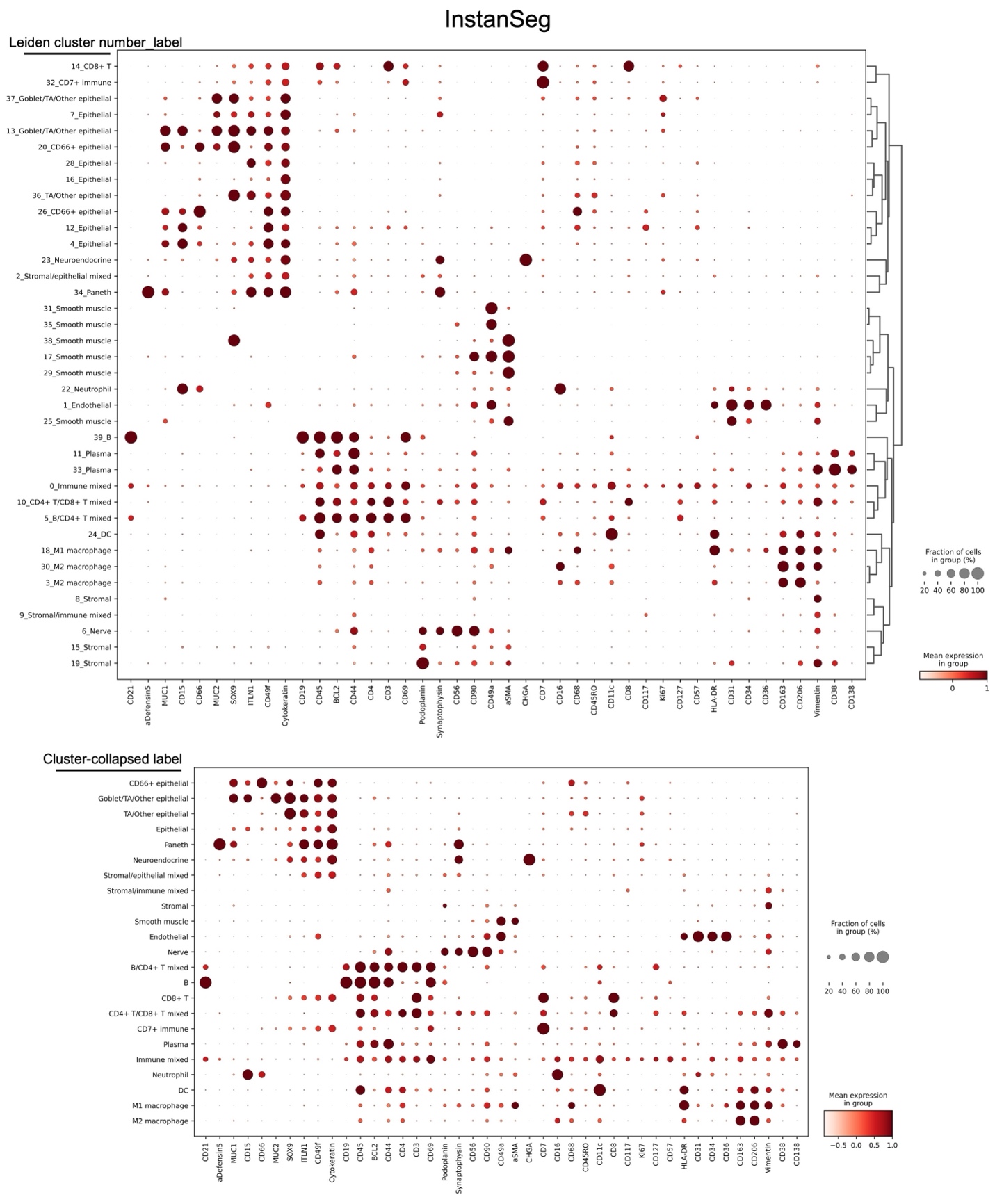

**Supplementary File 6. Leiden cluster labels before and after merging of identical labels for InstanSeg segmentation outputs.**

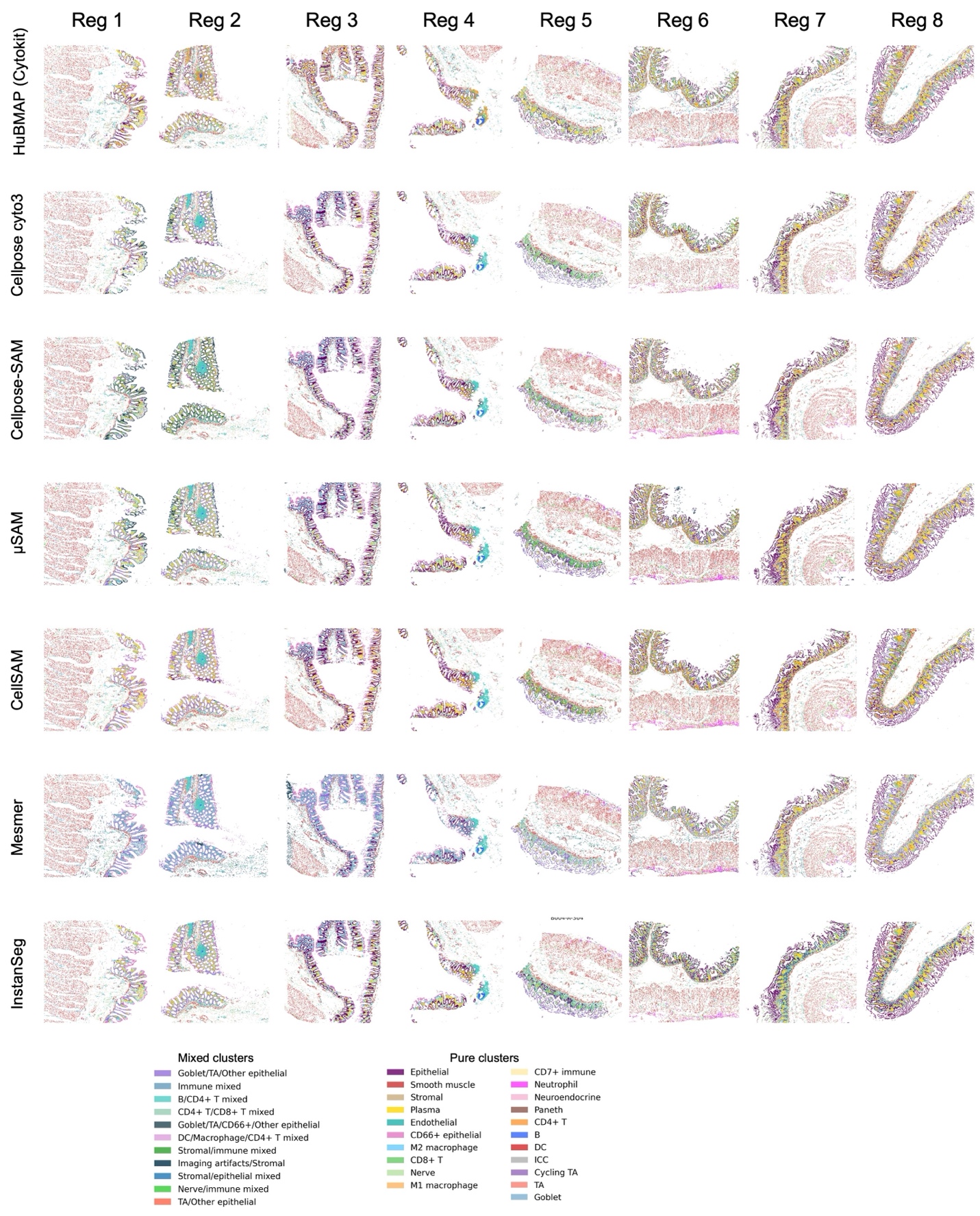

**Supplementary File 7. Spatial maps of all eight tissue regions across segmentation models on CODEX datasets after manual cell type annotation.**
